# Proteasome activator PA200 maintains stability of histone marks during transcription and aging

**DOI:** 10.1101/2020.04.28.067132

**Authors:** Tian-Xia Jiang, Shuang Ma, Xia Han, Zi-Yu Luo, Qian-Qian Zhu, Tomoki Chiba, Wei Xie, Su-Ren Chen, Kui Lin, Xiao-Bo Qiu

**Affiliations:** State Key Laboratory of Cognitive Neuroscience & Learning and Ministry of Education Key Laboratory of Cell Proliferation & Regulation Biology, College of Life Sciences, Beijing Normal University, 19 Xinjiekouwai Avenue, Beijing 100875, China; College of Life Sciences, Beijing Normal University, 19 Xinjiekouwai Avenue, Beijing 100875, China; Graduate School of Life and Environmental Sciences, University of Tsukuba, 1-1-1 Tennodai, Tsukuba, Ibaraki 305-8577, Japan; School of Life Sciences, Tsinghua University, Beijing 100084, China

**Author notes:** These authors contributed equally to this work (Tian-Xia Jiang, Shuang Ma, Xia Han). Correspondence (X.-B.Q.) or (K. L.).

**Keywords:** Histone marks, aging, histone degradation, PA200, proteasome, transcription

## Abstract

The epigenetic inheritance relies on stability of histone marks, but various diseases, including aging-related diseases, are usually associated with alterations of histone marks. How the stability of histone marks is maintained still remains unclear. The core histones can be degraded by the atypical proteasome, which contains the proteasome activator PA200, in an acetylation-dependent manner during somatic DNA damage response and spermiogenesis. Here we show that PA200 promotes the transcription-coupled degradation of the core histones, and plays an important role in maintaining the stability of histone marks. Degradation of the histone variant H3.3, which is incorporated into chromatin during transcription, was much faster than that of its canonical form H3.1, which is incorporated during DNA replication. This degradation of the core histones could be suppressed by the transcription inhibitor, the proteasome inhibitor or deletion of PA200. The histone deacetylase inhibitor accelerated the degradation rates of H3 in general, especially its variant H3.3, while the mutations of the putative acetyl-lysine-binding region of PA200 abolished histone degradation in the G1-arrested cells, supporting that acetylation is involved in the degradation of the core histones. Deletion of PA200 dramatically altered deposition of the active transcriptional hallmarks (H3K4me3 and H3K56ac) and transcription, especially during cellular aging. Furthermore, deletion of PA200 or its yeast ortholog Blm10 accelerated cellular aging. Notably, the PA200-deficient mice displayed a range of aging-related deteriorations, including immune malfunction, anxiety-like behaviors and shorter lifespan. Thus, the proteasome activator PA200 is critical to the maintenance of the stability of histone marks during transcription and aging.

## Introduction

The current model for epigenetic inheritance relies on stability of histone marks. The core histones, including H2A, H2B, H3 and H4, form an octamer to pack DNA into the nucleosome, the basic unit of chromatin (1). Post-translational modifications of histones (i.e., histone marks) regulate various cellular processes, including epigenetic regulation of transcription. For example, H3K4me3 and H3K56ac mark transcription initiation (2) and transcriptionally-active chromatin areas (3), respectively. Histone marks remain relatively stable in a specific type of cell or tissue, and should only be rearranged during development or cellular reprogramming (4, 5). However, aging and various diseases are usually associated with alterations of histone marks (6). Replacement of histone variants often occurs during transcription. Histone variant H3.3, which is expressed and incorporated into the chromatin in a replication-independent manner, is associated with transcriptional activation in higher eukaryotes (7). The transcription-coupled replacement of H3.1 with H3.3 occurs at gene bodies, promoters, and enhancers (8). H3.3 incorporation into nucleosomes marks the transcriptionally-active chromatin to control neuronal synaptic connectivity and cognition (9). It is a mystery how the stability of histone marks is maintained.

Proteasomes catalyze degradation of most cellular proteins, and consist of a 20S catalytic particle and one or two activators, such as the 19S regulatory particle, PA28α/β, PA28γ and PA200/PSME4 (10, 11). The typical 26S proteasome with the 19S particle as the activator promotes the ubiquitin-dependent protein degradation (12). PA200 and its yeast ortholog, Blm10, bind to the ends of the 20S particle (13, 14). PA200 accumulates on chromatin in response to DNA damage (15). We have recently shown that the core histones can be degraded by the PA200-proteasome in an acetylation-dependent manner during somatic DNA damage response and spermiogenesis (16). During elongation of spermatids, most core histones are degraded by the specialized PA200-containing proteasomes (i.e., spermatoproteasomes) (16). The acetylation-dependent histone degradation can also occur during DNA damage-induced replication stress (17). In response to DNA damage, the levels of histones from chromatin drop 20–40% in a manner depending on the INO80 nucleosome remodeler (18). In addition, histones are partially lost across the genome during aging in both yeast and human cells (19, 20). It remains unclear how histone loss or degradation influences the stability of histone marks.

In this study, we show that the core histones are degraded in the G1-arrested cells, and present evidence that PA200 promotes the transcription–coupled degradation of the core histones, and maintains the stability of histone marks during transcription and aging.

## Results

### PA200-proteasome promotes acetylation-dependent degradation of core histones in G1-arrested cells

To test whether histones are degraded during transcription, we took advantage of a modified pulse-chase assay by metabolically labeling proteins with a substitute of methionine (Met), azidohomoalanine (Aha). In this assay, Aha was co-translationally incorporated into proteins and subsequently ligated with biotin. Thus, following chase in the regular medium with Met, old histones with Aha could be purified with streptavidin for analysis of degradation (*SI Appendix,* Fig. S1*A*). In the G1-arrested mouse embryonic fibroblast (MEF) cells, the levels of histone H3 decreased during chase (Fig. 1A-C and *SI Appendix,* Fig. S1*B*). Histone variant H3.3, which can be incorporated into chromatin independently of replication, was also degraded in the G1-arrested MEF cells (Fig. 1A-C). Treatment with the proteasome inhibitor, Bortezomib (also named Velcade), blocked their degradation (Fig. 1A and B), suggesting that the proteasome mediates histone degradation in the G1-arrested MEF cells. Addition of Bortezomib increased the levels of histones at 0 h probably by blocking histone degradation, but decreased the levels of biotinylated histones probably by reducing the integration of Aha into proteins. Furthermore, the transcription inhibitor, α-amanitin, which induces degradation of the largest RNA polymerase II subunit Rbp1 (21), suppressed degradation of histones H3.3 and H3 (Fig. 1B). Because amanitin was added for 24 h before pulse chase, Rbp1 was not detectable even at 0 h of the chase. Because H3.3 is transcription-specific variant of H3, the levels of H3.3, but not of H3, decreased markedly at 0 h of the chase. These results strongly support that the degradation of the core histones depends on transcription.

**Fig. 1.**
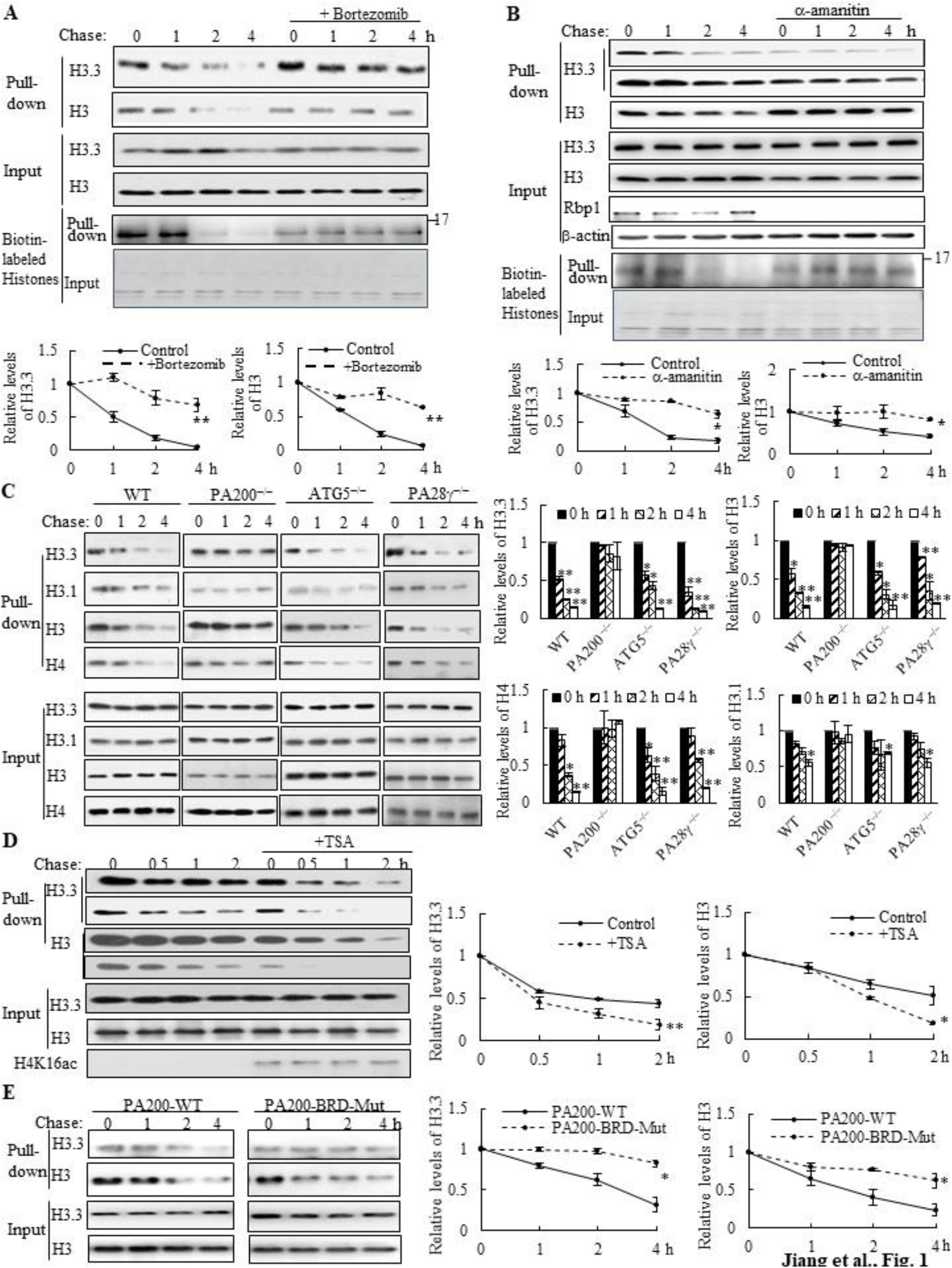
PA200 promotes acetylation-dependent degradation of core histones during transcription. (*A*) The G1-arrested MEF cells were pulse-labeled with Aha and then chased in the Met-containing medium in the absence or presence of 0.1 μM of Bortezomib for the time indicated. Bortezomib was added for 4 h at all time points of chases. Histones were captured by streptavidin-coupled beads and analyzed by immunoblotting. The corresponding input histone were used as the loading control. The captured histones levels were quantified by densitometry (normalized to the corresponding input histones). (*B*) Histone degradation in the G1-arrested MEFs treated with the transcription inhibitor α-amanitin at the concentration of 10 mM for 24 h was analyzed by the pulse-chase assay. The captured histones levels were quantified by densitometry (normalized to the corresponding input histone). (*C*) Histone degradation in the G1-arrested wild-type, PA200^−/−^, ATG5^−/−^, and PA28γ^−/−^ MEF cells was analyzed by the pulse-chase assay. Protein levels were analyzed by immunoblotting as in (*A*). (*D*) The G1-arrested MEF cells were incubated in the absence or presence of 0.3 μM of TSA for 4 h. Histone degradation was analyzed by the pulse-chase assay. Protein levels were quantified by densitometry (normalized to input histones). (*E*) Histone degradation in the G1-arrested 293T transfected with either the wild-type or mutant PA200 (PA200-BRD-Mut) with a HA-tag was analyzed by the pulse-chase assay. Protein levels were analyzed by immunoblotting as in (*A*). The relative histone levels in different groups at time point 0 were set to 1. All biotin-labeled histones in pull-down experiments were probed by the anti-streptavidin antibody, and the input biotin-labeled histones were analyzed by Coomassie blue staining in (*A-B*). Data are representative of one experiment with at least two independent biological replicates (mean ± SEM, ** *p* < 0.01, * *p* < 0.05, two-tailed unpaired *t*-test).

We previously showed that the PA200-proteasomes promote the acetylation-dependent degradation of the core histones during DNA repair and spermiogenesis (16). Deletion of PA200 suppressed the degradation of H3.3 and histone H4 in the G1-arrested cells (Fig. 1C). The canonical histone H3 variant H3.1, which is incorporated during replication (22), showed a slight degradation. This degradation was also suppressed by PA200 deletion (Fig. 1C and *SI Appendix,* Fig. S1*C*). However, deletion of another proteasomal activator, PA28γ, showed no inhibitory effects on histone degradation (Fig. 1C and *SI Appendix,* Fig. S1*C*), supporting the specific role of PA200 in histone degradation in the G1-arrested MEF cells. Lamin B1 is a nuclear protein, which undergoes the autophagy-mediated degradation (23). We found that the degradation of lamin B1 could not be blocked by deletion of PA200 or PA28γ (*SI Appendix,* Fig. S1*D*). A small fraction of histones is present in the cytosol and can be removed by macroautophagy, which requires the autophagic gene ATG5 (24). Our method using acid extraction could efficiently extract the core histones from the chromatin fraction to exclude the influence of the cytosolic fraction (25). Deletion of ATG5 had no effect on the degradation of chromatin histones, but suppressed degradation of lamin B1 (Fig. 1C and *SI Appendix,* Fig. S1*D*). Furthermore, trichostatin A (TSA), a histone deacetylase inhibitor, accelerated the degradation rates of H3 in general, especially of its variant H3.3 (Fig. 1D and *SI Appendix,* Fig. S1*E*). PA200 bears an acetyl-lysine-binding bromodomain-like (BRDL) region with critical residues at F2125 and N2126D (16). The mutations at this BRDL region (F2125S/N2126D) of PA200 abolished histone degradation in the G1-arrested cells (Fig. 1E and *SI Appendix,* Fig. S1*F*). The BRDL mutant had a more prominent effect on transcription-specific H3.3 than H3. These results further support our previous notion that acetylation is involved in the degradation of the core histones (16).

### PA200 promotes degradation of core histones primarily in actively-transcribed regions

By utilizing Aha to label proteins metabolically, we analyzed histone degradation genome-wide by sequencing the DNA fragments purified together with histones following 2 h-pulse labeling with Aha and chase in the regular medium for 0 or 4 h. In the wild-type MEF cells at 0-h chase, the levels of the Aha-labeled core histones (primarily H3 and H4) peaked at transcriptional start sites (TSS) genome-wide (Fig. 2A). In the wild-type MEF cells at 4-h chase, the levels of the Aha-labeled core histones dramatically dropped, but maintained at a similar pattern in coding regions to that at 0-h chase (Fig. 2A). In the PA200-deficient MEF cells at 0-h chase, the enrichment of the Aha-labeled core histones was much lower throughout the coding regions than that in the wild-type cells. In the PA200-deficient MEF cells at 4-h chase, the levels of the Aha-labeled core histones just slightly dropped in coding regions, supporting the notion that PA200 promotes degradation of the core histones during transcription. Surprisingly, the levels of the Aha-labeled core histones were raised at TSS in the PA200-deficient MEF cells after 4-h chase (Fig. 2A). Given that the nucleosome turnover by which the core histones are often disassembled from coding regions and then reassembled during transcription is exceptionally active at TSS (26), this rise is probably because deletion of PA200 led to accumulation of the Aha-labeled core histones at TSS regions in the process of nucleosome turnover. These results raise the possibility that the PA200-mediated degradation of the core histones facilitates nucleosome turnover.

**Fig. 2.**
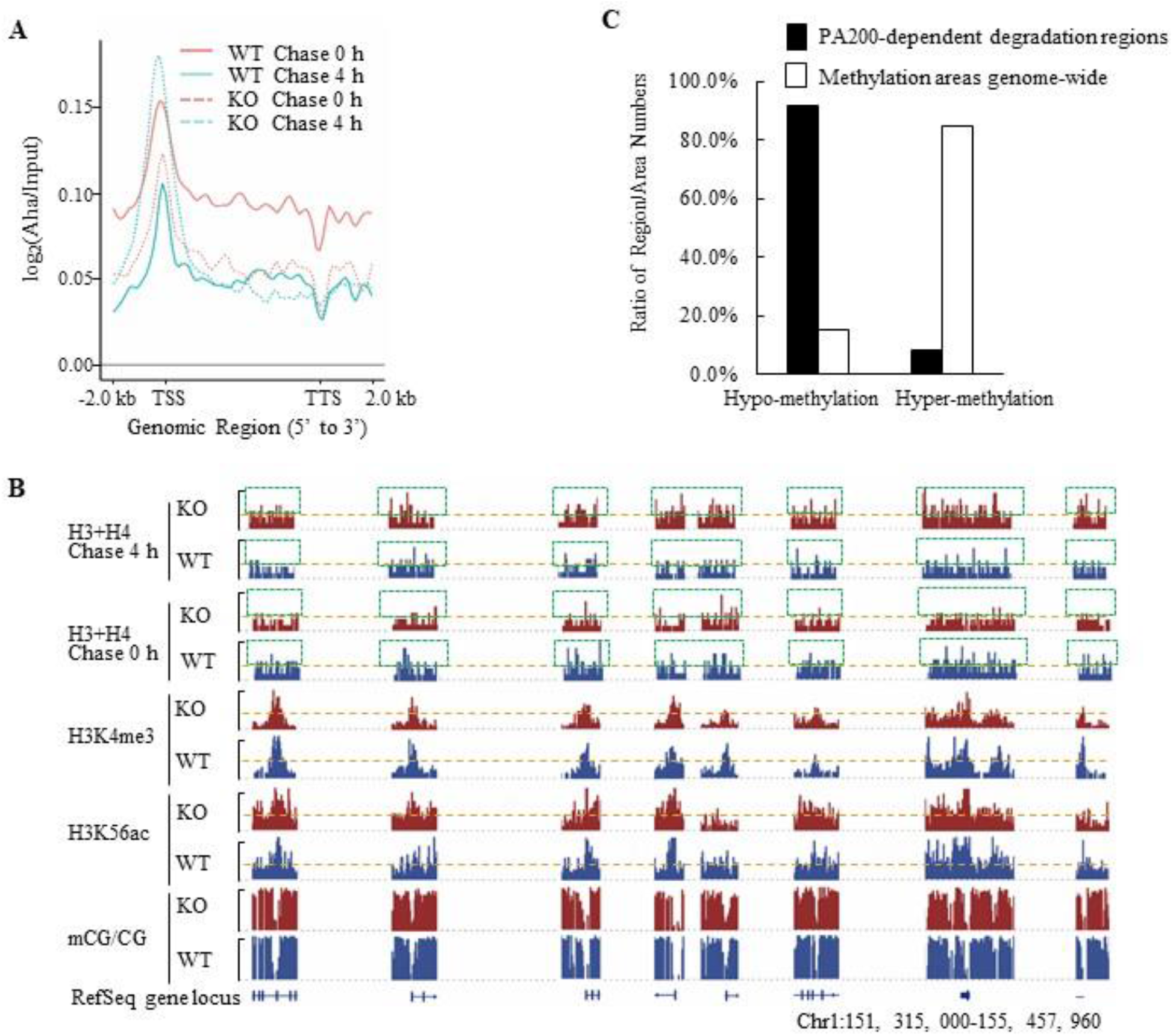
PA200 promotes degradation of core histones primarily in actively-transcribed regions. (*A*) Gene plot of core histone enrichment signals in PA200^+/+^ (WT) and PA200^−/−^ (KO) MEFs. (*B*) The IGV genome browser view of H3K4me3 and H3K56ac enrichment in WT and PA200 KO MEFs for the selected chromosomes, while other regions between them are hidden as indicated by grey dash lines. The levels of DNA methylation are shown in parallel. Yellow dash lines indicate equal heights at y axes in each pair of WT and PA200 KO samples. Squares with green dash lines highlighted the differences in the levels of histones H3 and H4 in the selected clusters of gene loci. The signal track of each sample was calculated by MACS2 and normalized by sequencing depth. (*C*) The distribution of PA200-dependent histone degradation regions genome-wide by collectively analyzing DNA sequencing and WGBS data. The PA200-dependent histone degradation regions include 5972 hypo-methylation regions and 550 hyper-methylation regions, which are defined in the Methods. The average CG levels within the “hyper-methylation” regions are >0.5. To analyze the distribution of hypo-/hyper-methylation areas genome-wide, the areas whose average CG levels are>0.5 are defined as “hyper-methylation areas”, whereas whose average CG levels are<0.5 are defined as “hypo-methylation areas” as described in the Methods. The numbers of hyper-methylation and hypo-methylation areas are 684 and 123, respectively.

The IGV genome browser view of these sequencing data further supported the above conclusions. Deletion of PA200 indeed reduced the enrichment of the Aha-labeled core histones in many genome regions, as evidenced by the reduced levels of the Aha-labeled core histones (e.g., areas highlighted by green squares) in the PA200-deficinet MEF cells at 0 h of chase (Fig. 2B). After 4-h chase, deletion of PA200 markedly attenuated the drop in the levels of the Aha-labeled core histones in most genome regions, and increased their levels (e.g., areas highlighted by green squares) at certain genome regions. H3K4me3 and H3K56ac mark transcription initiation (2) and transcriptionally-active chromatin regions (3), respectively. In general, there were only 8% (550/6522 regions) of the PA200-dependent degradation of the core histones was located in the hyper-methylation regions in the G1-arrested cells by analyzing DNA methylation data of whole-genome bisulfite sequencing (WGBS) along with the ChIP-seq data for histones (Fig. 2C). The ratio of the hyper-methylation areas genome-wide is 84.76% (684/807 areas) (Fig. 2C), suggesting that the core histones are primarily degraded in a PA200-dependent manner in the actively-transcribed regions (i.e., hypo-methylated regions). Taken together, these data demonstrate that the PA200-proteasome promotes the acetylation-dependent degradation of the core histones primarily in the actively-transcribed genome regions, and that degradation of the core histones is associated with the altered patterns of H3K4me3 and H3K56ac genome-wide.

### PA200 is critical to maintenance of histone marks and regulation of transcription

To investigate the role of PA200 in regulating transcription, the genome-wide RNA-sequencing analysis was performed using the total RNAs extracted from mouse livers and MEF cells, respectively. Significant changes in the expression of 334 genes were identified in the PA200-deficient livers (Fig. 3A and *SI Appendix,* Fig. S2*A* and *B*). Based on KEGG (Kyoto Encyclopedia of Genes and Genomes) database, functions of these differentially-expressed genes (DEGs) are mainly related to metabolism, transcription, signal transduction, cell growth, and cell death (*SI Appendix,* Fig. S2*C* and *D*). By analyzing the pathway enrichment of the RNA-seq data, we found that the metabolic pathways involved in the insulin signaling, gluconeogenesis and lipid metabolism were the top enriched pathways in the PA200-deficient livers (*SI Appendix,* Fig. S2*E*). Given that the liver plays critical roles in glucose/lipid metabolism, these results suggest that the PA200-dependent regulation of gene expression is critical to liver function.

**Fig. 3.**
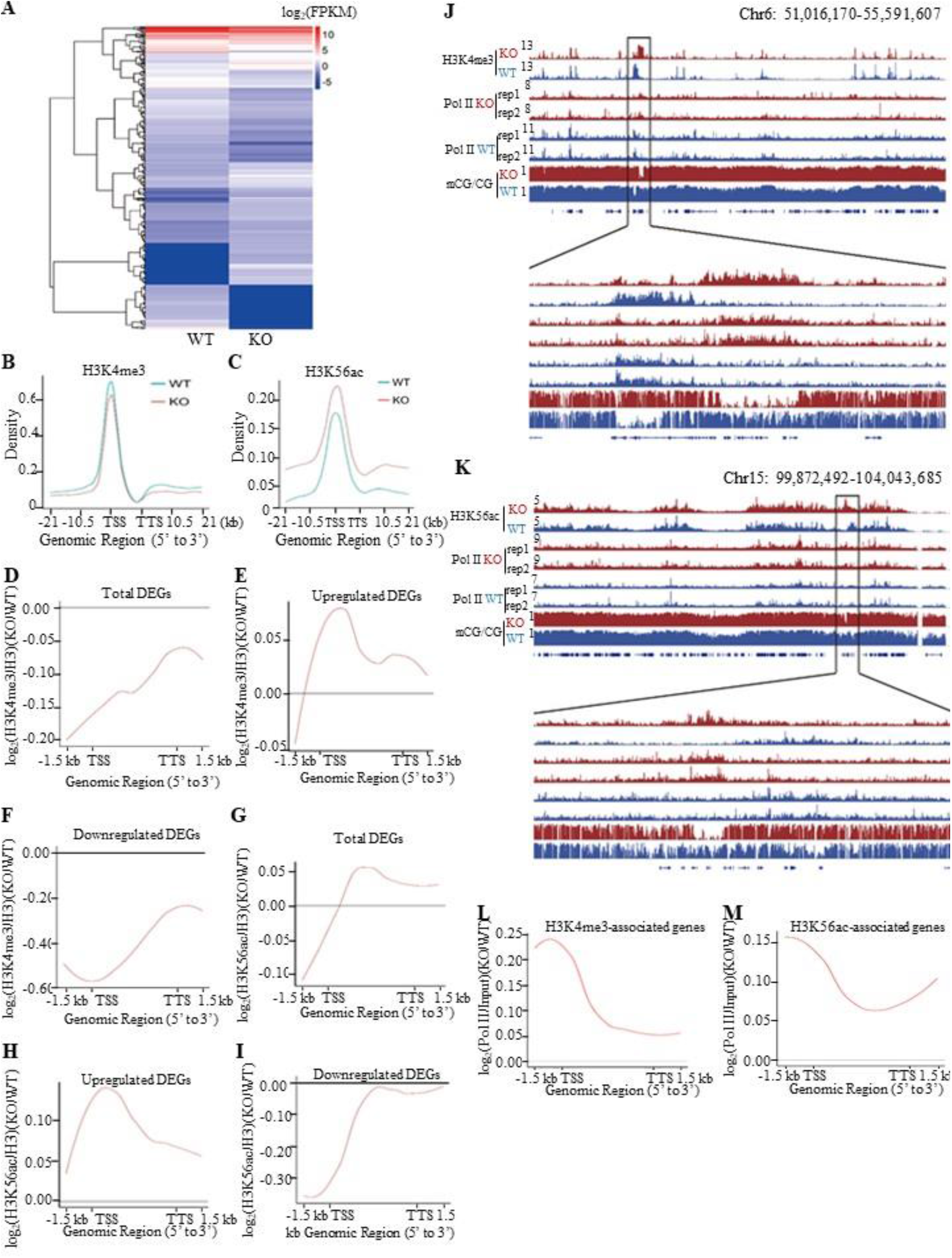
Deletion of PA200 disrupts genome-wide transcription. (*A*) Hierarchical clustering of the differentially-expressed genes (DEGs) in the PA200^+/+^ (WT) and PA200^−/−^ (KO) mouse liver. DEGs were defined according to the combination of the absolute value of log2-Ratio ≥1 and diverge probability≥0.8. Coloring indicates the log2-transformed fold change. (*B-C*) Enrichment of H3K4me3 (*B*) and H3K56ac (*C*) on the gene bodies and 21 kb upstream of TSS and 21kb downstream of TTS regions in the MEF genome. (*D-I*) The genome average plot for the change in H3K4me3 (*D-F*) or H3K56ac (*G-I*) (normalized to H3) in the PA200^−/−^ (KO) over the wild-type MEF cells (WT). (*J-K*) The IGV genome browser view of H3K4me3 (*J*) or H3K56ac (*K*) enrichment in PA200^+/+^ (WT) and PA200^−/−^ (KO) MEF chromosomes. The levels of DNA methylation and polymerase II were shown in parallel. (*L-M*) The profile analysis of RNA polymerase II on the gene bodies and 1.5 kb upstream of TSS and 1.5 kb downstream of TTS regions of the genes enriched with the H3K4me3 (*L*) or H3K56ac (*M*) in the PA200^−/−^ (KO) MEF cells normalized to those in the wild-type group (WT).

Accordingly, deficiency of PA200 in the G1-arrested MEF cells markedly influenced the transcription of 1314 genes (*SI Appendix,* Fig. S3*A-C*), which regulate important pathophysiological processes, such as aging, metabolism and cancer (*SI Appendix,* Fig. S3*D-F*). Only a few genes were co-up or down-regulated in the PA200-deficient livers and the G1-arrested PA200^−/−^ MEF cells (*SI Appendix,* Fig. S3*G*). Actually, the coverage of the aging pathways is incomplete in these databases. Thus, we searched literatures for the aging-related DEGs of RNA-seq in mouse livers and MEFs, respectively, and the results showed that deletion of PA200 caused the up- or down-regulation of almost completely different sets of the aging-related genes (*SI Appendix,* Table. S1). Taken together, the PA200-mediated proteolysis regulates transcription of genes, which dictate many critical pathophysiological activities, at both cell and tissue levels.

To explore the mechanism underlying the above transcriptional changes, we performed chromatin-immunoprecipitation (ChIP)-sequencing analyses of histone marks, which revealed a slight down-regulation of H3K4me3 on the gene bodies and 21 kb upstream of TSS and 21 kb downstream of TTS regions in the G1-arrested PA200^−/−^ MEF cells (Fig. 3B). In contrast, deletion of PA200 led to a marked upregulation of H3K56ac in these genome regions (Fig. 3C). Consistently, PA200 deletion markedly elevated protein levels of H3K56ac, but slightly increased the levels of H3K4me3, as revealed by immunoblotting assays (*SI Appendix,* Fig. S4*A*). The up-regulated DEGs were accompanied with the increased H3K4me3 and H3K56ac marks, while the down-regulated DEGs were with the reduced H3K4me3 and H3K56ac marks in the G1-arrested PA200-deficient MEF cells (Fig. 3D-I, and *SI Appendix,* Fig. S4*B* and *C*). In contrast, neither the up-regulated DEGs (including *Hoxa9, Ctnna2,* and *Bmp4*) nor the down-regulated DEGs (including *Fzd2, SOD1,* and *HMGB1*) showed these effects in the PA200-deficient MEF cells that were not arrested (*SI Appendix,* Fig. S4*B* and *C*). KEGG analysis of the DEGs associated with H3K4me3 or H3K56ac revealed their involvements in the cell cycle, senescence, autophagy, and the cancer- or development-related signal pathways (*SI Appendix,* Fig. S5*A-D*). H3K4me3 and H3K56ac showed similar regulatory tendency on certain up-regulated or down-regulated genes, but displayed distinct tendency on most genes, in response to PA200 deficiency in MEF cells (*SI Appendix,* Fig. S5*E* and *F*). It is noteworthy that the number for the upregulated genes associated with H3K4me3 or H3K56ac enrichment was 2-3 folds of that for the downregulated genes, hinting an indirect role for PA200 in suppressing expression of most genes because both H3K4me3 and H3K56ac are usually associated with active transcription (27). Notably, changes in the deposition of H3K4me3 or H3K56ac were positively correlated with the recruitment of RNA polymerase II onto chromatins, and inversely correlated with DNA methylation in certain critical gene regions (Fig. 3J and K, and *SI Appendix,* Fig. S6*A-H*). Furthermore, the recruitment profile of RNA polymerase II resembled that for the upregulated DEGs in both H3K4me3- and H3K56ac-associated genes in the PA200-deficient MEF cells as shown above (Fig. 3E, H, L, and M), suggesting that PA200 regulates transcription by acting at the upstream of RNA polymerase II recruitment.

Deletion of PA200 increased the levels of DNA methylation in general (*SI Appendix,* Fig. S7*A-C*), changed distributions of differential methylation regions (DMRs) (*SI Appendix,* Fig. S7*D* and *E*), and influenced diverse cellular functions (*SI Appendix,* Fig. S7*F*). Although there were certain overlaps between changes in DMRs and the differentially-expressed genes in PA200^−/−^ MEFs (*SI Appendix,* Fig. S7*G* and *H*), most hypo-DMR-related genes were upregulated, while a large portion of hyper-DMR-related genes were downregulated *SI Appendix,* Fig. S7*G-J*). Moreover, the inverse correlations of H3K4me3 or H3K56ac with DMR were primarily observed within gene regions (*SI Appendix,* Fig. S7*K-O*), and a certain portion of this inversion corresponded to the genes that were upregulated (*SI Appendix,* Fig. S7*P* and *Q*). Taken together, these results suggest that PA200 is critical to the maintenance of histone marks in gene regions and to the regulation of transcription.

### PA200 regulates transcription of aging-related genes

To analyze the nature of the genes affected by PA200 deletion, expressions of all genes were grouped into 2 clusters (up- and down-regulated). Enrichment of H3K4me3 or H3K56ac at promoters was generally in a positive correlation with the changes in the RNA levels of specific genes (Fig. 4A). Pathway enrichment analysis of DEGs revealed that PA200 deficiency changed the expression of many aging-related genes, such as aging-promoting genes: *Bmp4* (28), *Cdkn1a* (*p21*^*Cip1*^) (29), *Cdkn2b* (*p15*^*INK4b*^) (30) and aging-suppressing genes: *Hmgb1* (31) and *Sod1* (32) (Fig. 4B and Table S1). The H3K4me3 levels and the H3K56ac levels markedly increased at gene bodies in the up-regulated aging-related genes (Fig. 4C and D, and *SI Appendix,* Table S1). The levels of both H3K4me3 and H3K56ac were reduced in the down-regulated aging-related genes in the PA200-deficient MEF cells (Fig. 4E and F, and *SI Appendix,* Table S1). Specifically, H3K56ac and H3K4me3 were enriched on the gene bodies of certain aging-promoting genes, such as *Cdkn2a* (*p16*^*INK4a*^) (33) and *Bmp4*, and decreased on gene bodies of some aging-suppressing genes, such as *Hmgb1* and *Sirt2* (34) in PA200^−/−^ MEF cells (Fig. 4G-N). Quantitative PCR analyses confirmed that expression of these aging-related genes was altered accordingly in the PA200-deficient mouse liver (*SI Appendix,* Fig. S8*A*). These results suggest that PA200 regulates the transcription of aging-related genes.

**Fig. 4.**
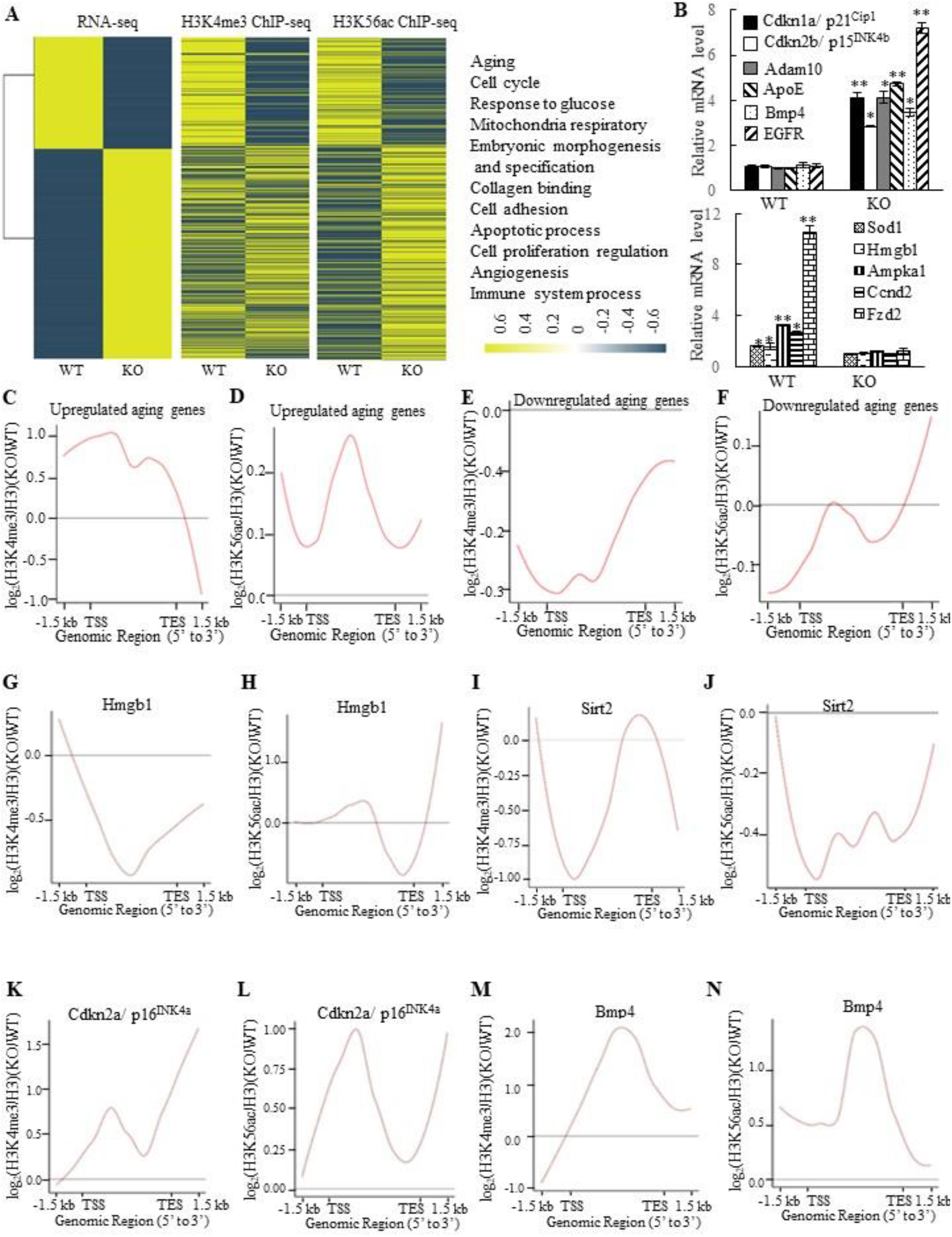
Deletion of PA200 alters transcription of the aging-related genes. (*A*) Heat-map comparison of all DEGs with the H3K4me3 or H3K56ac enrichment at promoters (TSS ± 2.5 kb; middle) in the PA200-deficient MEF cells (normalized to the wild-type group). DEGs were defined according to the combination of the absolute value of log2-ratio ≥1 and diverge probability≥0.8. Coloring indicates the log2-transformed fold change. DEGs were clustered into 11 major groups with enriched GO terms listed (right). All *p* values of GO Terms are <0.0001. (*B*) Quantitative PCR analysis of expression of the aging-related genes in MEF cells. (*C-F*) The genome average plot for the change in H3K4me3 (*C, E*) or H3K56ac (*D, F*) (normalized to H3) on up-regulated or down-regulated aging-related genes in the PA200^−/−^ (KO) normalized to the wild-type group. (*G-N*) The profile analysis of H3K4me3 (*G, I, K, M*) and H3K56ac (*H, J, L, N*) on the gene bodies of *hmgb1, sirt2, p16*^*INK4a*^, and *bmp4* in the PA200^−/−^ (KO) MEF cells normalized to the wild-type group. Data are representative of one experiment with at least three independent biological replicates (mean ± SEM, ** *p* < 0.01, * *p* < 0.05, two-tailed unpaired *t*-test).

### PA200/Blm10 extends cellular lifespan

We next found that deletion of PA200 reduced the proliferative potential (*SI Appendix,* Fig. S8*B*), and increased the activity of the senescence-associated β-galactosidase (SA-β-gal), a marker for cellular aging (35), in primary MEF cells (Fig. 5A). Significant changes in the expression of 3740 genes were identified in the PA200-deficent primary MEFs at days 0 and 30 in comparison to those in the wild-type cells. Meanwhile, these changes in DEGs displayed distinct patterns of expression between day 0 and day 30 (Fig. 5B). Deletion of PA200 disrupted the expression of the genes related to aging and other pathophysiological processes, including metabolism, development, cancer, and genetic information processing (*SI Appendix,* Fig. S8*C-G*). The levels of H4 usually decrease, but the levels of H4K16ac increase, during aging (19). Notably, deletion of PA200, but not of PA28γ suppressed the loss of H4 during aging (Fig. 5C). In order to examine whether this role of PA200 is conserved evolutionally, we showed that deletion of the PA200 ortholog Blm10 decreased, but overexpression of Blm10 increased, the yeast bud scar numbers (Fig. 5D), which represent replicative lifespan (19). Overexpression of Blm10 accelerated, but deletion of Blm10 prevented, the decrease in the levels of both H4 and H4K16ac during aging (Fig. 5E and F, and *SI Appendix,* Fig. S9), supporting the notion that Blm10 promotes the acetylation-mediated histone degradation during aging. These results suggest that PA200/Blm10 regulates transcription during cellular aging and extends cellular lifespan.

**Fig. 5.**
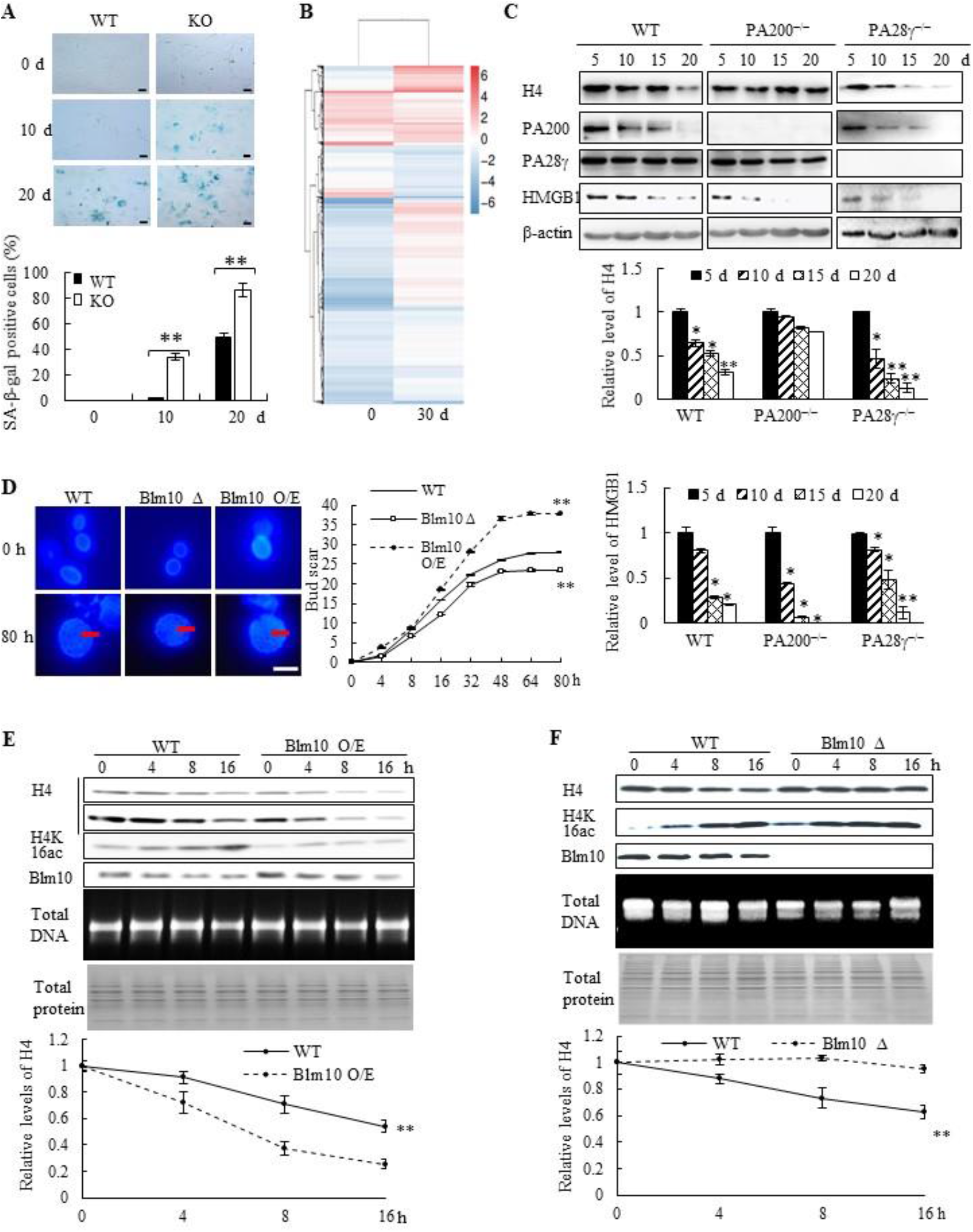
PA200/Blm10 promotes degradation of the core histones during aging and extends cellular lifespan. (*A*) SA-β-gal staining of primary WT and PA200^−/−^ (KO) MEF cells at the indicated days. The percentage of β-galactosidase-positive cells in each group was displayed in bar graph. Scale bar: 50 μm. (*B*) Heat map showing the expression (normalized reads per kilo base per million mapped-reads (RPKM)) of all DEGs in PA200^−/−^ (KO) MEF cells cultured for 0 or 30 days (normalized to WT MEFs). Each treatment has two independent biological replicates which were clustered into one group. Coloring indicates the log2 transformed fold change. (*C*) Immunoblotting of the whole-cell extracts from primary MEF cells at indicated days. The levels of HMGB1, a biomarker for young cells, decreased during aging. Protein levels were quantified by densitometry (normalized to β-actin). (*D*) Bud scars of the wild-type, Blm10-deficient (Blm10Δ) or Blm10-overexpressing (Blm10 O/E) yeast at the indicated times. The bar graph showed the bud scar numbers, and images from the calcofluor 28 staining showed the sizes of yeast cells and bud scars. A scar is pointed by an arrow. More than 30 cells were counted for each group. (*E*) Immunoblotting of the whole-cell extracts of the wild-type or Blm10 O/E at the indicated times. Total sonicated DNA on 1% agarose gel was used as control. The total proteins were visualized by Coomassie staining following SDS-PAGE. Quantitation of histone levels was carried out by densitometry (normalized to total DNA). (*F*) Immunoblotting of the whole-cell extracts of the wild-type or Blm10Δ yeast as analyzed similarly to (*E*). Quantitation of histone levels was carried out by densitometry (normalized to total DNA). The relative histone levels in different groups at time point 0 were set to 1. Data are representative of one experiment with two independent biological replicates (mean ± SEM, ** *p*<0.01, * *p*<0.05, two-tailed unpaired *t*-test).

### PA200 deficiency accelerates aging-related pathological changes

To further explore whether PA200 participates in regulation of aging, we analyzed the wild-type and PA200-deficient mice for distinct parameters of aging. The aging-related increase in memory T cells and reduction in naive T cells are important predictors of senescence (36). At 12-month old, female PA200^−/−^ mice had more memory and fewer naive CD4T cells, however, these differences were not significant in the younger cohort (Fig. 6A). Renal glomerulosclerosis usually occurs with aging (37). We analyzed interstitial inflammation and glomerular hypercellularity for glomerulosclerosis in the PA200-deficient mice. The results showed that the ratio of sclerotic glomeruli markedly increased at 3- or 12-month old in the PA200-deficient mice in comparison to the wild-type mice (Fig. 6B). We also examined muscle loss, a hallmark of aging in both humans and rodents (38). Gastrocnemius muscle fiber diameter declined with age more seriously in PA200^−/−^ mice than that in the wild-type mice. Unlike the wild-type mice, which showed an 8% decrease in gastrocnemius muscle fiber diameter between 3 and 12 months old, the PA200-deficient mice had a 15% decrease with age (Fig. 6C), suggesting that PA200 deletion promotes muscle fiber atrophy. We then assessed anxiety-like behaviors in 12-month-old mice. PA200-deficient mice spent more time and made more errors than the wild-type mice (Fig. 6D and E), indicating that PA200 deficiency caused spatial learning defects in aging processes. Finally, we examined mouse lifespan, and found that deletion of PA200 reduced mouse lifespan (Fig. 6F). Taken together, these results suggest that deletion of PA200 leads to the aging-related deteriorations in mice.

**Fig. 6.**
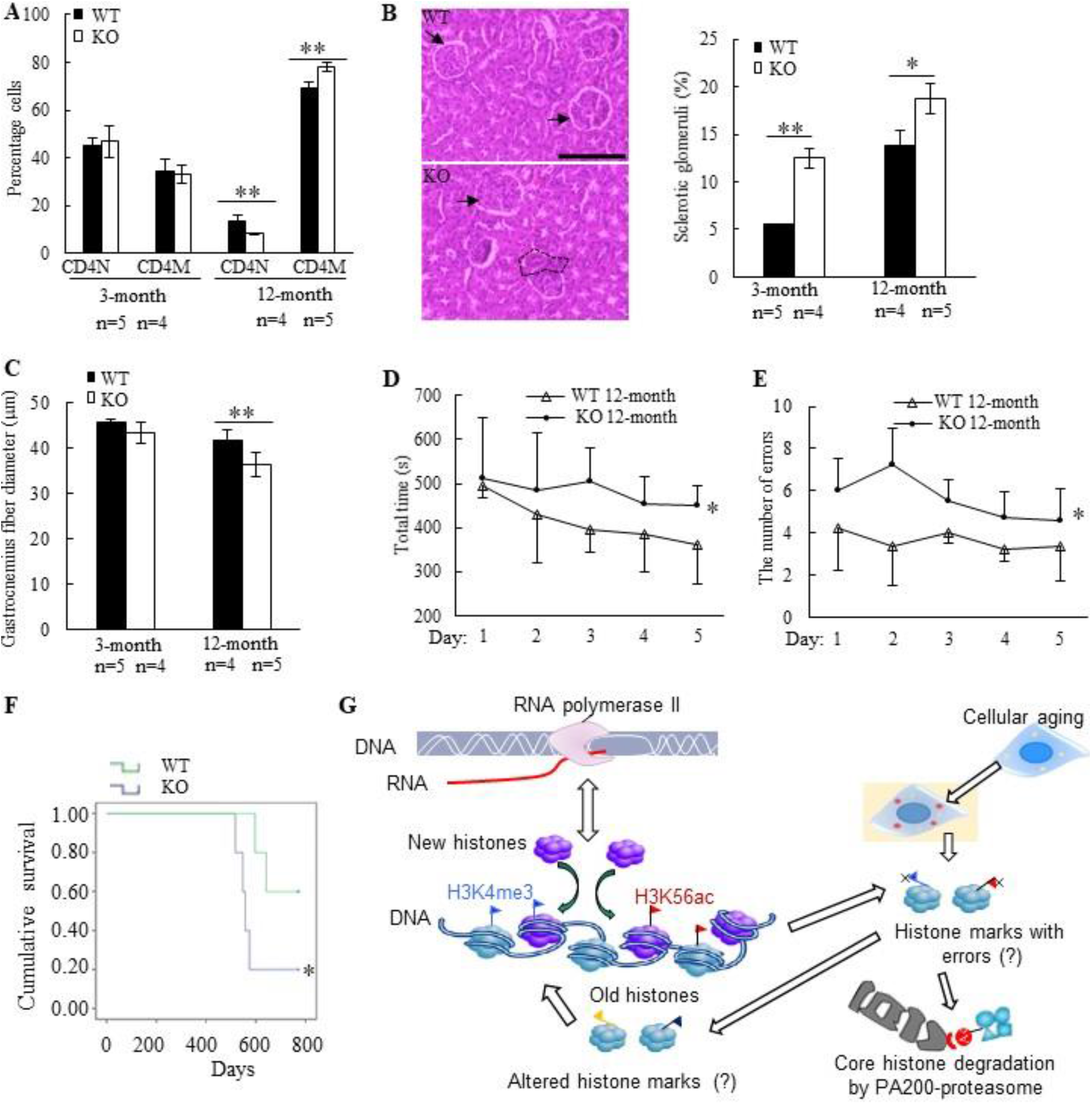
Deletion of PA200 accelerates aging in mice. (*A*) T cell subset analysis for naive CD4 (CD4N) and memory CD4 (CD4M) subsets in 3- and 12-month-old wild-type (WT) or PA200^−/−^ (KO) mice. (*B*) Hematoxylin-eosin stained kidney sections of 3- and 12-month-old mice. The phenotypes of renal sclerosis include interstitial inflammation (dashed area) and glomerular hypercellularity (as denoted by arrows) in KO mice (lower). The arrows denoted normal glomeruli in the wild-type mice (upper). Scale bar, 100 μm. The bar graph showed the percentage of sclerotic glomeruli from 12-month-old kidney sections. 40 glomeruli were scored for each animal. (*C*) The bar graph showed the gastrocnemius muscle fiber diameter of 3 and 12 months of age in WT and KO mice. (*D*) The time in accomplishing eight successful performances of 8-arm maze test by 12-month-old WT or KO mice. (*E*) The number of errors made before eight successful performances in 8-arm maze test. Wild-type group: n=4, PA200^−/−^ group: n=5. (*F*) Lifespan in WT (n = 5) and PA200 KO (n = 5) mice. (*G*) A schematic showing a hypothetical model for the mechanism by which PA200 promotes the acetylation-dependent proteasomal degradation of the core histones during transcription and aging, and regulates deposition of the active transcriptional hallmarks, such as H3K4me3 and H3K56ac, and transcription, probably by degrading histones with abnormal marks (Data are shown as mean ± SEM, ** *p*<0.01, * *p*<0.05, two-tailed unpaired *t*-test).

## Discussion

The pattern of the histone post-translational modifications has been proposed as “histone code” (39). As the potential carriers of epigenetic code, histones have long been believed to be non-degradable during transcription. Indeed, parental histones can pass into the next generation of cells during DNA replication (40). A previous study has implicated the involvement of the ubiquitin-mediated protein degradation by the proteasome in the nucleosome turnover (9). Since the ubiquitin-proteasome pathway might regulate the nucleosome turnover indirectly, *e.g.*, by degrading histone chaperones that assist assembly of the nucleosome, the transcription-coupled degradation of histones has not yet been validated. We have previously shown that the PA200-containing proteasomes can promote the acetylation-dependent degradation of the core histones during somatic DNA repair and spermatogenesis (16). Moreover, Mandemaker et al. showed that the PA200-proteasome is required for the acetylation-dependent degradation of the core histones during DNA damage-induced replication stress (17). This study demonstrated that the PA200-containing proteasome promoted degradation of the core histones during transcription. Degradation of the histone variant H3.3, which is incorporated into chromatin during transcription, was much faster than that of its canonical form H3.1, which is incorporated during replication (22). As a support, we showed that the core histones were degraded in the G1-arrested cells. This degradation of the core histones could be suppressed by the transcription inhibitor, the proteasome inhibitor or deletion of PA200, but not by deletion of another proteasome activator PA28γ. Notably, the histone deacetylase inhibitor accelerated the degradation rates of H3 in general, especially its variant H3.3, while the mutations of the putative acetyl-lysine-binding region of PA200, which specifically binds the acetylated core histones *in vitro* (16), abolished histone degradation in the G1-arrested cells. This is consistent with our previous notion that acetylation is involved in degradation of the core histones (16). Thus, our results suggest that the core histones are degradable during transcription, and PA200 promotes their degradation in an acetylation-dependent manner.

The 19S complex of the 26S proteasome was shown earlier to function independently of the 20S catalytic particle, playing a direct and non-proteolytic role in RNA polymerase II-mediated transcription (41). But, it was suggested recently that the transcriptionally-relevant form of the proteasome is the canonical 26S complex (42), hinting that the degradation of certain proteins by the 26S proteasome might influence transcription indirectly. We have shown previously that the 19S particle is not directly involved in the degradation of the core histones, which were ectopically expressed (16). The situation could be different during transcription. Even during spermiogenesis, deletion of PA200 just retards the degradation of the core histones at step 11 of spermatogenesis, and the core histones are eventually degraded in the survived elongated spermatids and sperm in the PA200-deficient mice (16). As a result, the PA200-deficient mice are still fertile, though with the dramatically reduced fertility (43). This study demonstrates that PA200 regulates deposition of the transcriptionally-active histone marks, including H3K4me3 and H3K56ac. This deposition was positively correlated with the recruitment of RNA polymerase II onto chromatins, and inversely correlated with DNA methylation, which usually marks transcriptionally-inactive region in certain critical gene regions (44). Thus, these results further suggest that PA200 assures the deposition of proper histone marks.

Partial histone loss across the genome has been shown to be associated with aging in both yeast and human cells (19, 20). Transcription of various genes is altered during aging (45–47). The aging-related changes in transcription have been suggested to be caused by histone loss in yeast (8). This study showed that both PA200 and its yeast ortholog Blm10 promoted proteasomal degradation of the core histones during aging, and regulated transcription during cellular aging. Furthermore, deletion of either PA200 or Blm10 accelerated cellular aging. Notably, the PA200-deficient mice displayed a range of aging-related deteriorations, including immune malfunction, anxiety-like behaviors, and the reduced lifespan.

Although the cellular aging can prevent tumor development early in life, aging is also marked by an increase in tumorigenesis (45, 48, 49). Both aging and tumorigenesis are accompanied with the accumulation of abnormal histone marks (50). PA200 deficiency caused abnormal transcription of many genes involved in cancers and altered deposition of H3K4me3 or H3K56ac on the genes related to cancer signaling, such as those in the p53, MAPK, and Ras signaling pathways. We speculate that these aging-related phenotypes of the PA200-deficient mice might be a consequence of the accumulated “old” histones with abnormal histone marks.

It has been a mystery how histone marks remain stable during transcription. Histones are frequently evicted from the nucleosome and then re-assembled in a transcription-coupled manner, a process referred to as nucleosome turnover (26). Histone exchange is generally highest at active promoters, where H3K4me3, H3K56ac, H3K9ac and H3K14ac usually accumulate (51). It can be imagined that certain mistakes in reassembling must happen during transcriptional nucleosome turnover, especially in the aged cells. Thus, the PA200/Blm10-mediated degradation of the core histones would probably eliminate the abnormally-assembled histone marks during transcription (Fig. 6G). Our results also raise the possibility that the PA200-mediated degradation of the core histones facilitates nucleosome turnover. Although we could not exclude the possible involvement of other substrates of PA200 or an indirect role of PA200, the acetylation-dependent degradation of the core histones must play an important role in these PA200-related activities. In conclusion, our results suggest that PA200 maintains the stability of histone marks during transcription and aging.

## Materials and Methods

Detailed methods are available in SI Appendix, SI Methods.

### Strains and Cell Culture

MEF cells were cultured in Dulbecco’s modified Eagle’s medium (DMEM) supplemented with 10% fetal bovine serum (FBS), 100 U/ml penicillin, 100 μg/ml streptomycin, 1% non-essential amino acids and 200 μM of β-mercaptoethanol. Primary PA200^+/+^, PA200^−/−^ MEF cells, and PA28γ^−/−^ MEF cells were isolated from the embryos of the wild-type, PA200-deficient mice and PA28γ-deficient mice, respectively. The permanent wild-type and PA200-deficient MEF cells were obtained from the mice as described (16). The PA28γ-deficient cells were obtained from Drs. Lance Barton and Xiaotao Li. The ATG5-deficient MEF cells were obtained from Dr. Alfred L. Goldberg, who received the cells from Dr. Yoshinori Ohsumi.

All yeast strains were grown at 30°C in yeast peptone dextrose (YPD). The yeast strain BY4741 (MATa his3△1 leu2△0 met15△0 ura3△0) was from Dr. Wei Li. The strain YHS539 (MATa BY4741 yak1-GFP (kan), blm10::Nat hat9) was obtained from Dr. Daniel Finley. The yeast strain Blm10 O/E (MATa BY 4741 blm10::NAT GPD-HA-Blm10) was obtained as described (16).

### Pulse-chase assay

The pulse-chase assay was employed to determine histone degradation by labeling proteins with Aha (52). Briefly, the G1-arested MEF cells cultured in MEF medium with minosine (0.2 μM, Sigma-Aldrich, #M0253) were first subjected to starvation with the media containing all amino acid except 0.2 mM methionine (Met) for 30 min. Then, the medium was replaced with MEF medium containing Aha (0.2 mM, Anaspec, #63669), instead of Met, for 2 h. After Aha treatment, the MEF medium with 0.2 mM Met was used to replace the medium with Aha. Because macroautophagy can remove the cytosolic fraction of the core histones (24), we used acid extraction, which could efficiently extract the core histones from the chromatin fraction, to exclude the cytosolic fraction of the core histones (25), and then ligated chromatin histones with biotin through the copper-catalyzed cycloaddion reaction. The Aha-containing histones of chromatin were captured by streptavidin-coated beads, and were finally incubated with 1× SDS buffer at 95 °C for 5 min. The samples were analyzed by immunoblotting to detect the corresponding proteins.

### Genome-wide analysis of histone degradation

After pulse-chase analysis as described as above, the DNA fragments associated with the newly-synthesized H3/H4 proteins were then purified with DNA purification columns (CST, #14209) and processed for ChIP-seq as described later.

We used MAnorm to statistically compare the signal density of ChIP-seq data at chase times 0 and 4 h. To identify the degraded region of the G1-arrested cells, we selected the unique peaks which were present in ChIP-seq at 0 h, but were absent at 4 h. We selected the common peaks whose read density at 0 h was significantly larger than that at 4 h. Then, we defined the PA200-depdentdent histone degradation regions, which were not overlapped between the wild-type and the PA200^−/−^ samples.

### Yeast replicative life span analysis

Old yeast cells were isolated by using EZ Link Sulfo-NHL-LC-LC-Biotin (Thermo Scientific) to label cell surface proteins (Invitrogen) (53). The older cells were isolated by several rounds of this affinity purification. Ages of isolated cells were estimated by counting Calcofluor-stained bud scars as described previously (19). Proteins on the surface of logarithmic phase cells were labeled with EZ Link Sulfo-NHL-LC-LC-Biotin. The old mother cells were isolated by Dynabeads Streptavidin T1, and stained with calcofluor 28 (Sigma) for counting the bud scar numbers.

### Immunoblotting assay

Cells were lysed in the buffer containing 20 mM Tris–HCl, pH 8.0, 100 mM KCl, 0.2% Nonidet P-40, 10% glycerol, 1mM ZnCl2, 10 mM β-glycerophosphate, 5 mM tetrasodium pyrophosphate, 1 mM NaF, 1 mM Na3VO4, and a mixture of protease inhibitors. Protein samples were separated by SDS-PAGE and transferred onto PVDF membranes. Antibodies or antisera against specific proteins were used as primary antibodies to detect the corresponding proteins. Peroxidase-conjugated anti-mouse IgG (1:5000) or anti-rabbit IgG (1:5000) was used as the secondary antibody. The protein bands were visualized by ECL detection system (Millipore) or ODYSSEY (LI-COR).

## Data availability

All sequencing data, including RNA–seq, ChIP–seq, genome-wide histone degradation sequencing, and WGBS, have been deposited at the BioProject database (https://www.ncbi.nlm.nih.gov/bioproject) with accession number PRJNA451247. The data that support the findings of this study are available from the authors upon request. Correspondence and requests for materials should be addressed to X.B.Q. or K.L.

## ACKNOWLEDGMENTS

We greatly appreciate Drs. Lance Barton, Daniel Finley, Alfred L. Goldberg, Wei Li, Xiaotao Li, and Yoshinori Ohsumi for kindly providing cell lines or strains. This study was supported by the National Key R & D Program of China (2019YFA0802100), National Natural Science Foundation of China (31530014), Ministry of Science and Technology of China (2018YFC1003300) and Beijing Municipal Natural Science Foundation (5202014).

## AUTHOR CONTRIBUTIONS

T.X.J. devised and performed all the experiments in mammalian systems, analyzed data and wrote the manuscript. S.M. devised and performed the experiments in yeast. Q.Q.Z., T.C. and Z.Y.L. assisted the construction of the PA200-deficient mice and mouse experiments. X.H., W.X. and K.L. analyzed data. X.B.Q. conceived the project, supervised the experiments, analyzed data, and wrote the manuscript.

## THE AUTHORS DECLARE NO CONFLICT OF INTEREST

## Supplementary Information

### Supplemental Experimental Results

**Fig. S1.**
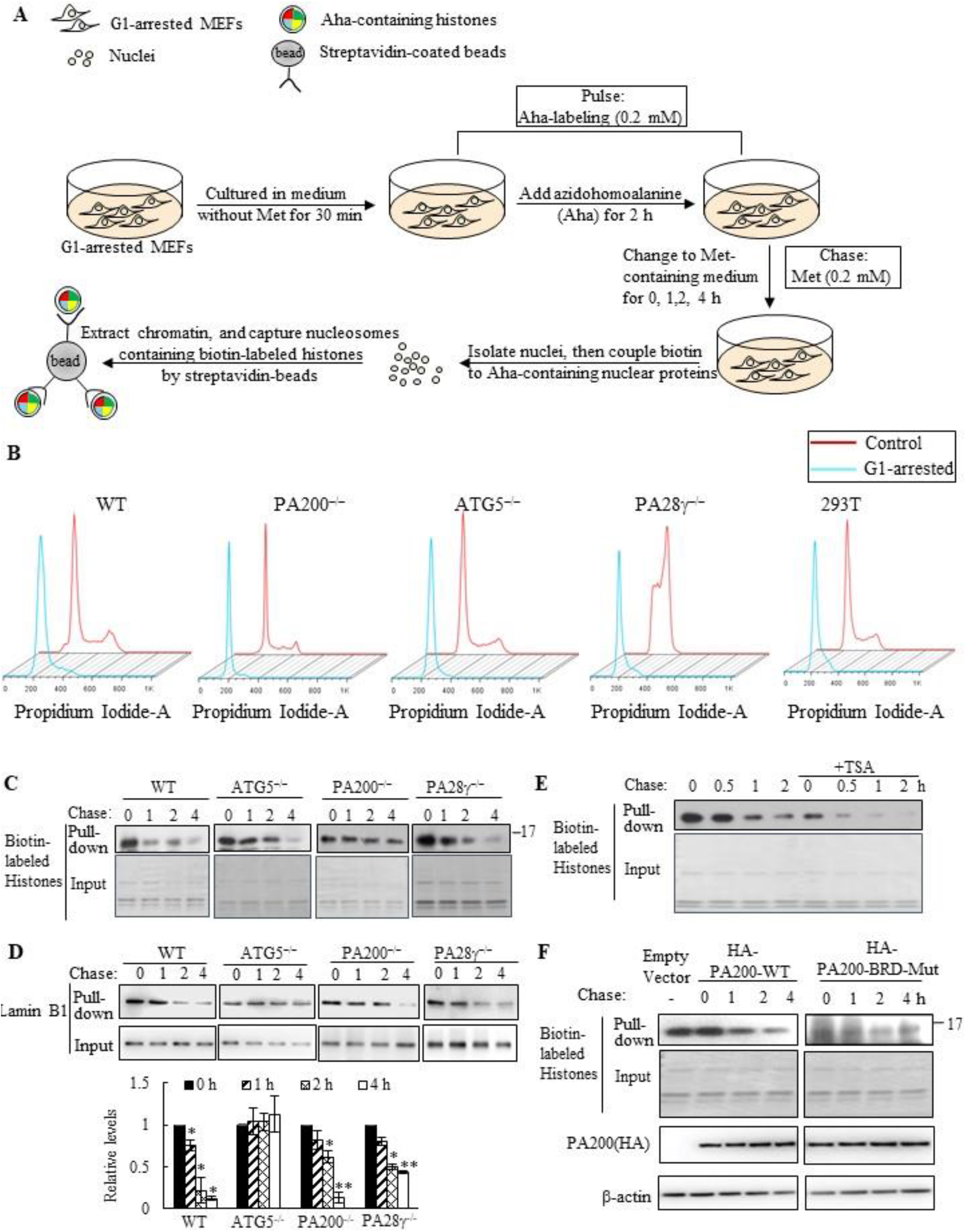
PA200 is essential for proteasomal degradation of core histones in G1-arrested cells. (*A*) Modified pulse-chase analysis of histone degradation during transcription. Aha was co-translationally incorporated into proteins and subsequently ligated with biotin. Following chase in the regular medium with Met, old histones with Aha was affinity-purified with streptavidin for analysis of degradation. (*B*) MEF cells cultured for 24 h in the basal medium were treated with or without 0.5 mM minosine for 24 h. DNA content was analyzed by FACS analysis of cellular DNA with propidium iodide staining. 10000 cells were collected for analyzing in each group. (*C, D*) Aha-labeled histone (*C*) or lamin B1 (*D*) was captured by streptavidin-coupled beads and analyzed with streptavidin-HRP or anti-lamin B1 antibody by immunoblotting in the G1-arrested wild-type, PA200^−/−^, ATG5^−/−^, and PA28^−/−^ MEF cells following the pulse-chase assay. The captured lamin B1 levels were quantified by densitometry (normalized to the corresponding input). (*E*) The G1-arrested MEF cells were incubated in the absence or presence of 0.3 μM of TSA for 4 h, and histone degradation was analyzed as described in (*C*). (*F*) Histone degradation in the G1-arrested 293T transfected with either wild-type or mutant PA200 (PA200-BRD-Mut) with a HA-tag was analyzed as described in (*C*). The transfection efficiency was confirmed by detecting the levels of HA-tag. The biotin-labeled input histones in (*E-F*) were analyzed by Coomassie blue staining. Data are representative of one experiment with at least two independent biological replicates (mean ± SEM, \*\**p* < 0.01,* *p* < 0.05, two tailed unpaired test).

**Fig. S2.**
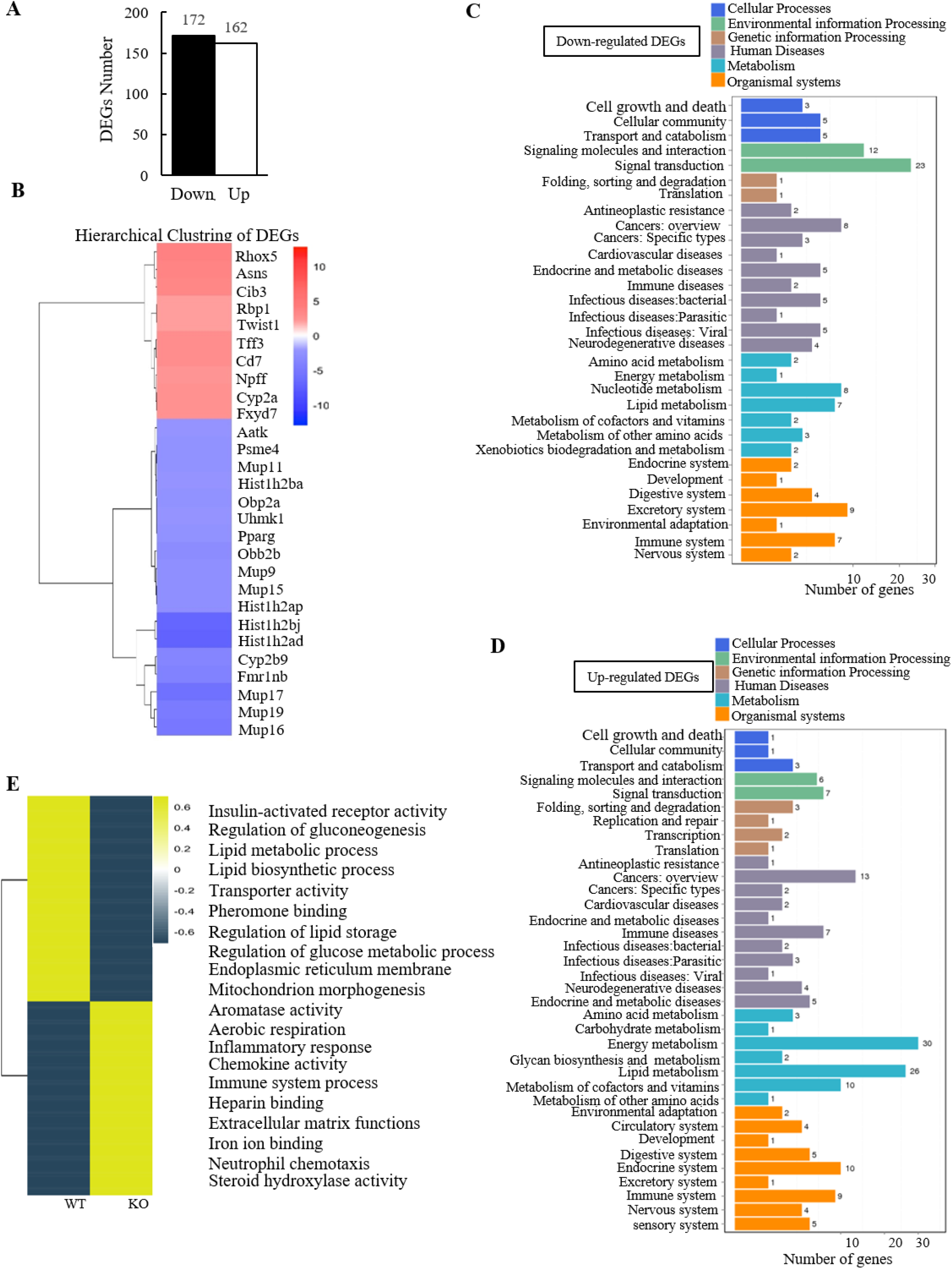
PA200 deletion influences gene expression as profiled by RNA-seq in mouse liver. (*A*) Numbers of the up-regulated and down-regulated genes in the PA200-deficient mouse liver (normalized to the wild-type group). (*B*) Hierarchical clustering of 28 selected DEGs. All selected DEGs are up-regulated two to four folds or down-regulated two to seven folds in PA200-deficient mouse liver relative to wild-type liver. (*C, D*) KEGG pathway classification of the down-regulated (*C*) and up-regulated (*D*) DEGs following RNA-seq analyses of the PA200-deficient mouse liver in comparison to those of the wild-type liver. (*E*) Heat-map of all DEGs in the PA200-deficient mouse livers. DEGs were defined according to the combination of the absolute value of log2-ratio ≥1 and diverge probability≥0.8. Coloring indicates the log2-transformed fold change. DEGs were clustered into 20 major groups with enriched GO terms listed (right). All *p* values of GO Terms are <0.05.

**Fig. S3.**
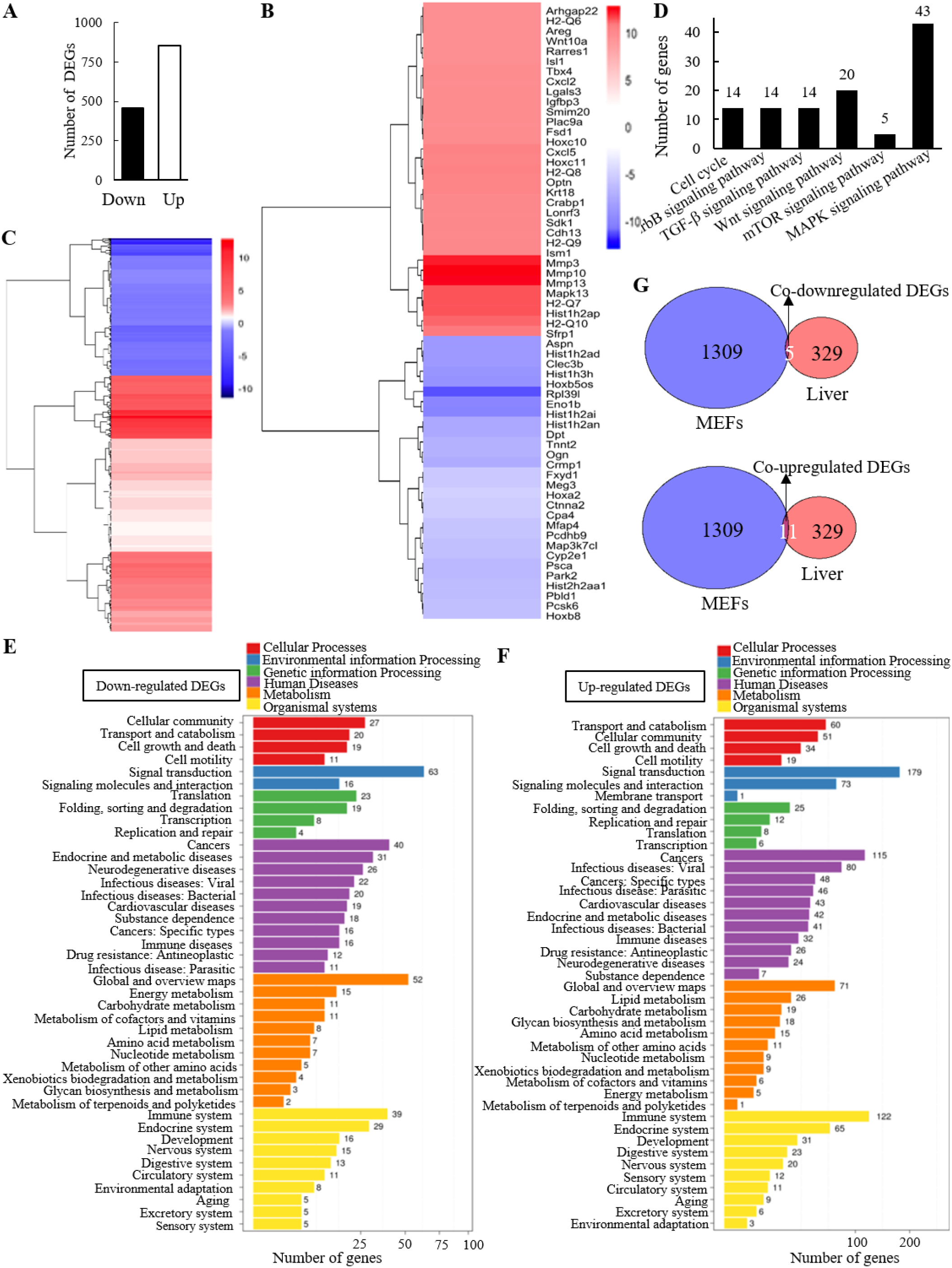
Gene expression profiling by RNA-seq in wild-type (WT) and PA200^−/−^ (KO) MEFs. (*A*) Numbers of the up-regulated and down-regulated genes in the PA200-deficient MEF cells (normalized to the wild-type group). (*B*) Hierarchical clustering of 61 selected DEGs. All selected DEGs are up-regulated ten to thirteen folds or down-regulated five to twelve folds in PA200^−/−^ MEFs relative to WT MEFs. (*C*) Hierarchical clustering of intersection DEGs in the PA200-deficient MEF cells (normalized to the wild-type group). DEGs were defined according to the combination of the absolute value of log2-Ratio ≥1 and diverge probability≥0.8. Coloring indicates the log2 transformed fold change. (*D*) KEGG classification into the aging-related pathways on DEGs in MEF cells. (*E, F*) KEGG pathway classification of down-regulated (*E*) and up-regulated (*F*) DEGs analyzed by RNA-seq in the PA200-deficient MEF cells (normalized to the wild-type group). (*G*) Co-up- or down-regulated genes in both the PA200-deficient livers and the G1-arrested PA200-deficient MEF cells.

**Fig. S4.**
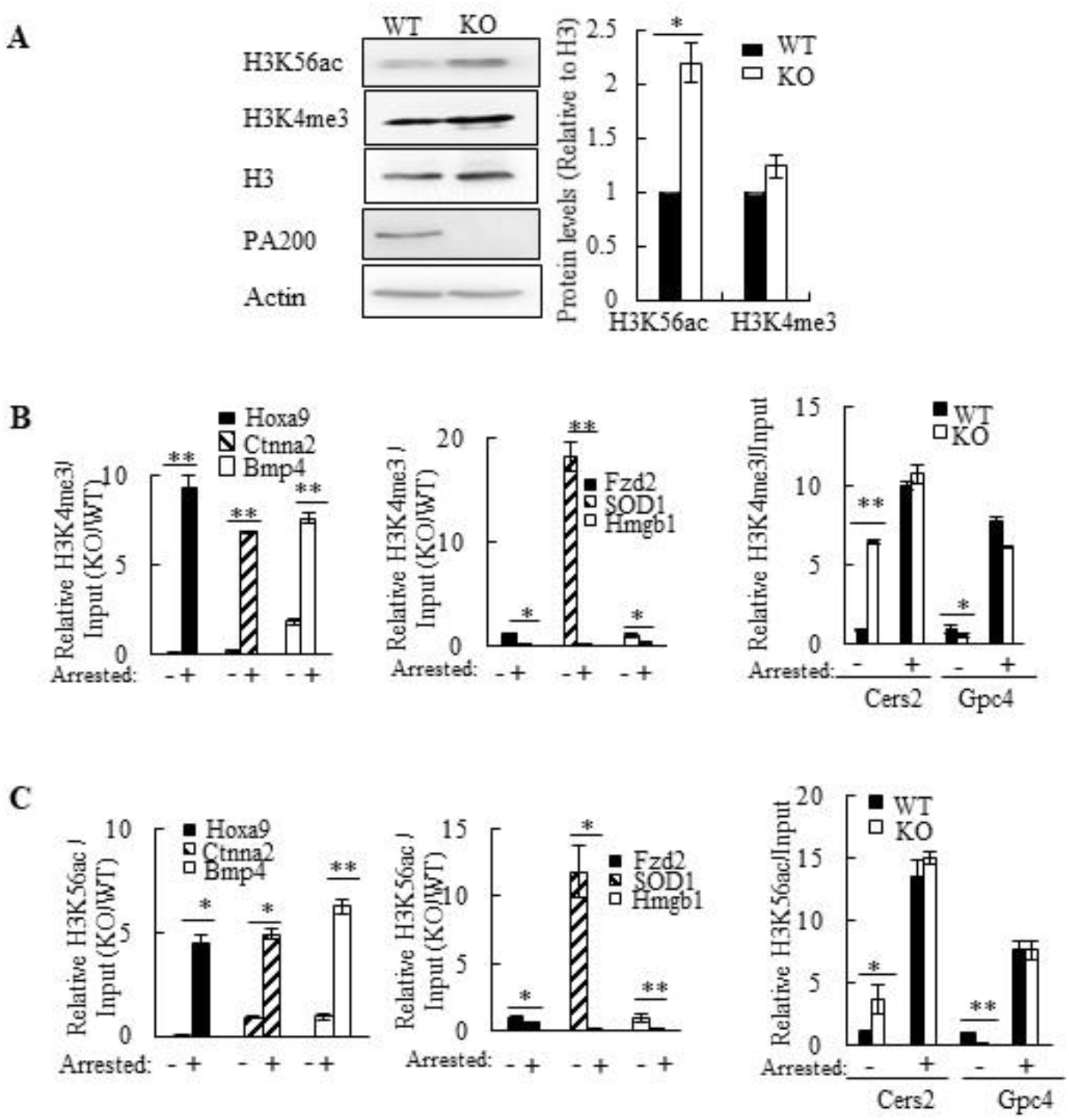
PA200 deletion affects protein levels of H3K56ac and H3K4me3, as well as the enrichments of the DEGs. (*A*) Immunoblotting analysis of H3K56ac or H3K4me3 levels in the PA200^+/+^ (WT) and PA200^−/−^ (KO) MEF cells. The levels of H3K56ac and H3K4me3 were quantified by densitometry (normalized to H3). (*B*) ChIP-PCR analysis of the promoter levels of the up-regulated DEGs (*Hoxa9, Ctnna2, Bmp4*), down-regulated DEGs (*Fzd2, SOD1, HMGB1*), and not significantly changed genes (*Cers2* and *Gpc4*) on H3K4me3 in the G1-arrested or non-arrested MEF cells. (*C*) ChIP-PCR analysis of the promoter levels of the up-regulated DEGs (*Hoxa9, Ctnna2, Bmp4*), down-regulated DEGs (*Fzd2, SOD1, HMGB1*), and not significantly changed genes (*Cers2 and Gpc4*) on H3K56ac in the G1-arrested or non-arrested MEF cells. Data are representative of one experiment with at least three independent biological replicates (mean ± SEM, ** *p* < 0.01, * *p* < 0.05, two tailed unpaired test).

**Fig. S5.**
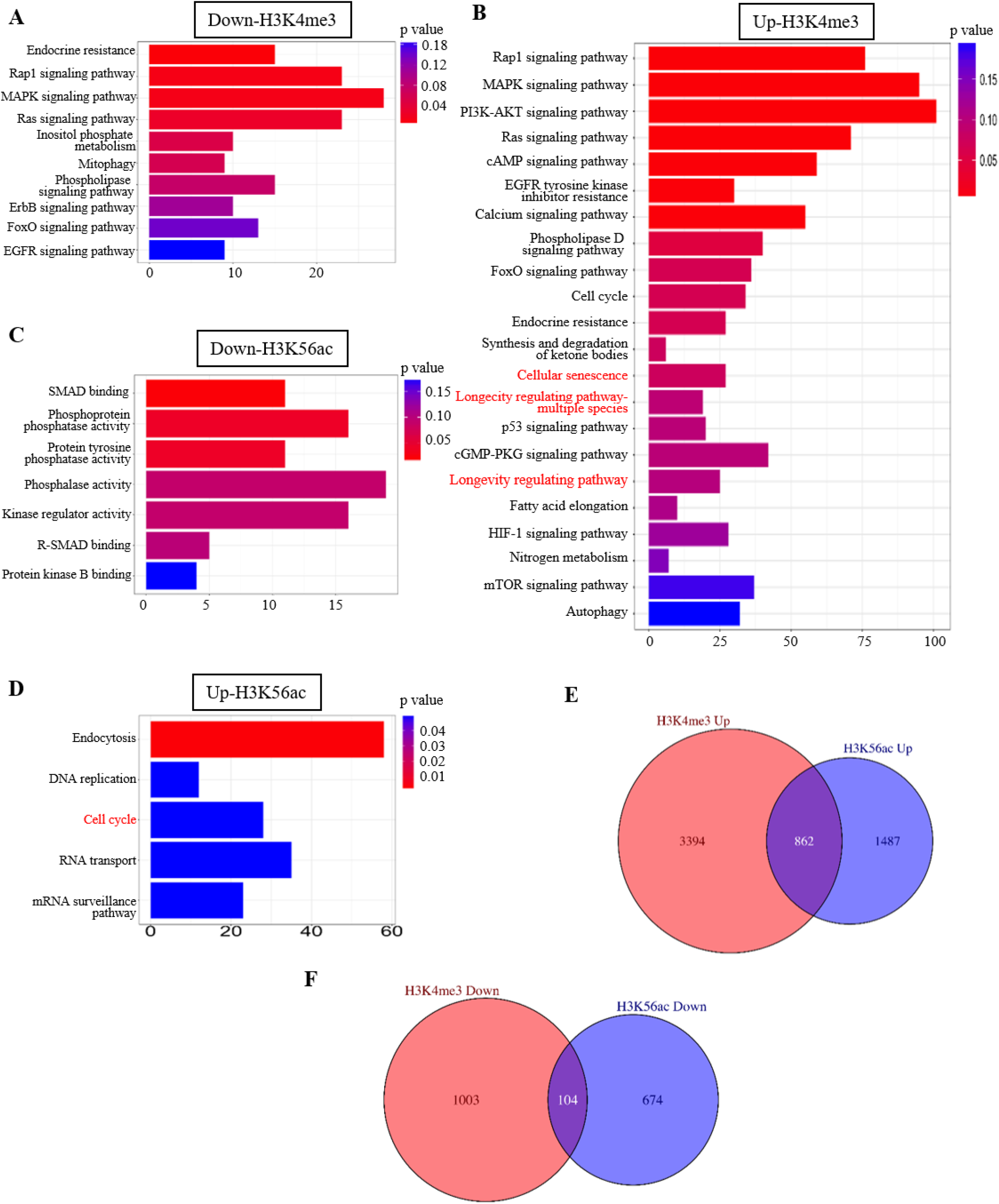
Analysis of H3K4me3 and H3K56ac-associated genes. (*A, B*) KEGG analysis of down-regulated (*A*) and up-regulated (*B*) regions-associated genes of H3K4me3 following ChIP-seq in the PA200-deficient MEF (normalized to the wild-type group). (*C, D*) KEGG analysis of genes within down-regulated (*C*) and up-regulated (*D*) H3K56ac-enriched regions in the PA200-deficient MEF (normalized to the wild-type group). If their levels of H3K4me3 or H3K56ac in PA200^−/−^ MEFs are higher than those in wild-type MEFs, these regions are defined as “Up-regulated” regions”. (*E, F*) Venn plots showing numbers of the common up-regulated (*E*) or down-regulated (*F*) genes between H3K4me3 and H3K56ac differential peaks in the PA200-deficient MEF cells (normalized to the wild-type group).

**Fig. S6.**
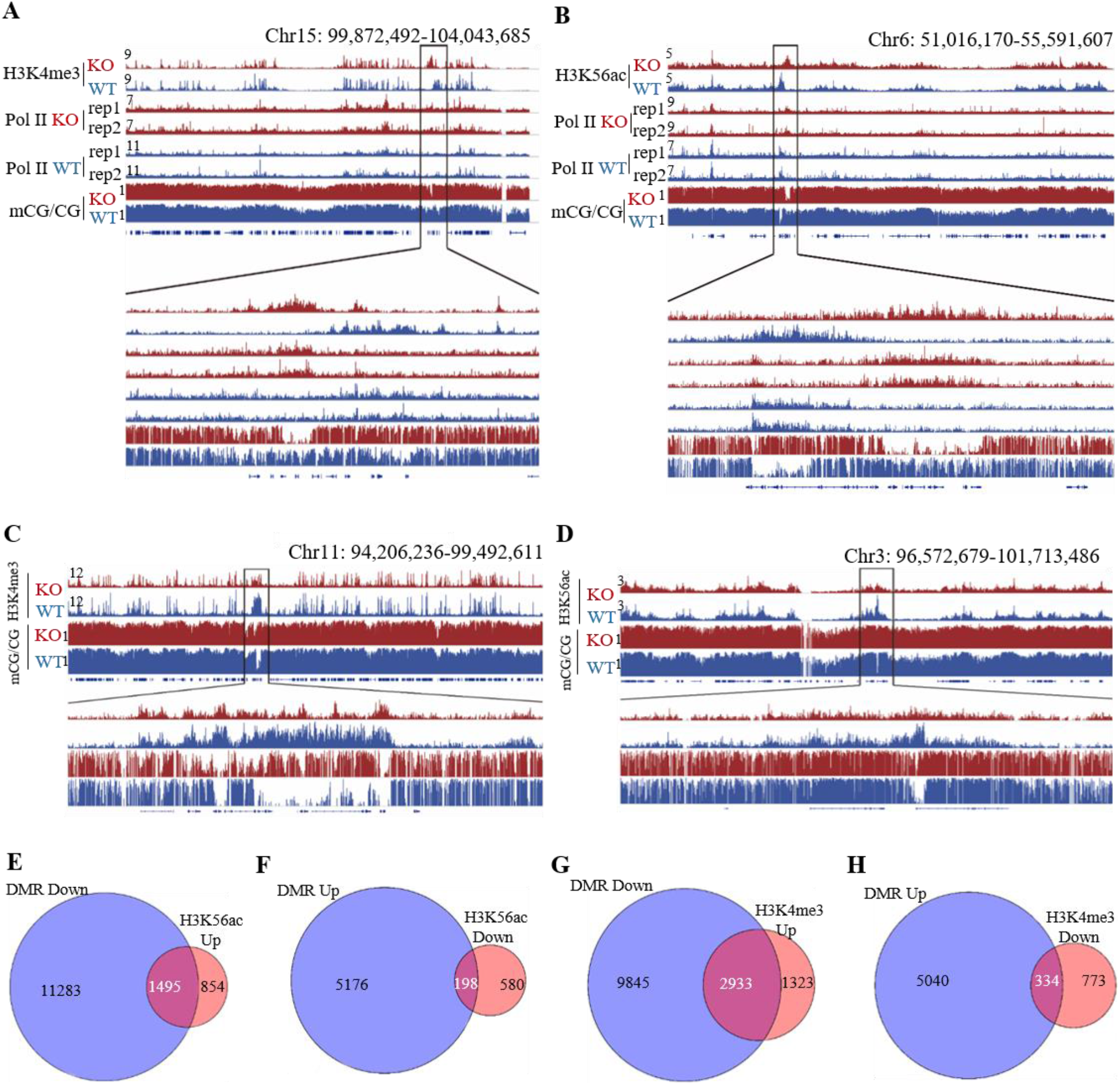
Changes of H3K56ac and H3K4me3 enrichments positively correlate with polymerase II recruitments, and inversely correlate with DNA methylation in certain gene regions. (*A-D*) The IGV genome browser view of H3K4me3 (*A, C*) or H3K56ac (*B, D*) enrichment in WT and PA200^−/−^ MEF chromosomes. The levels of DNA methylation and polymerase II are shown in parallel. (*E, F*) Venn plots showing numbers of genes that were up-regulated (*E*) or down-regulated (*F*) in H3K56ac differential peaks and inversely correlated with DNA methylation in certain gene regions in the PA200-deficient MEF cells. (*G, H*) Venn plots showing numbers of genes that were up-regulated (*G*) or down-regulated (*H*) in H3K4me3 differential peaks and inversely correlated with DNA methylation in certain gene regions in the PA200-deficient MEF cells.

**Fig. S7.**
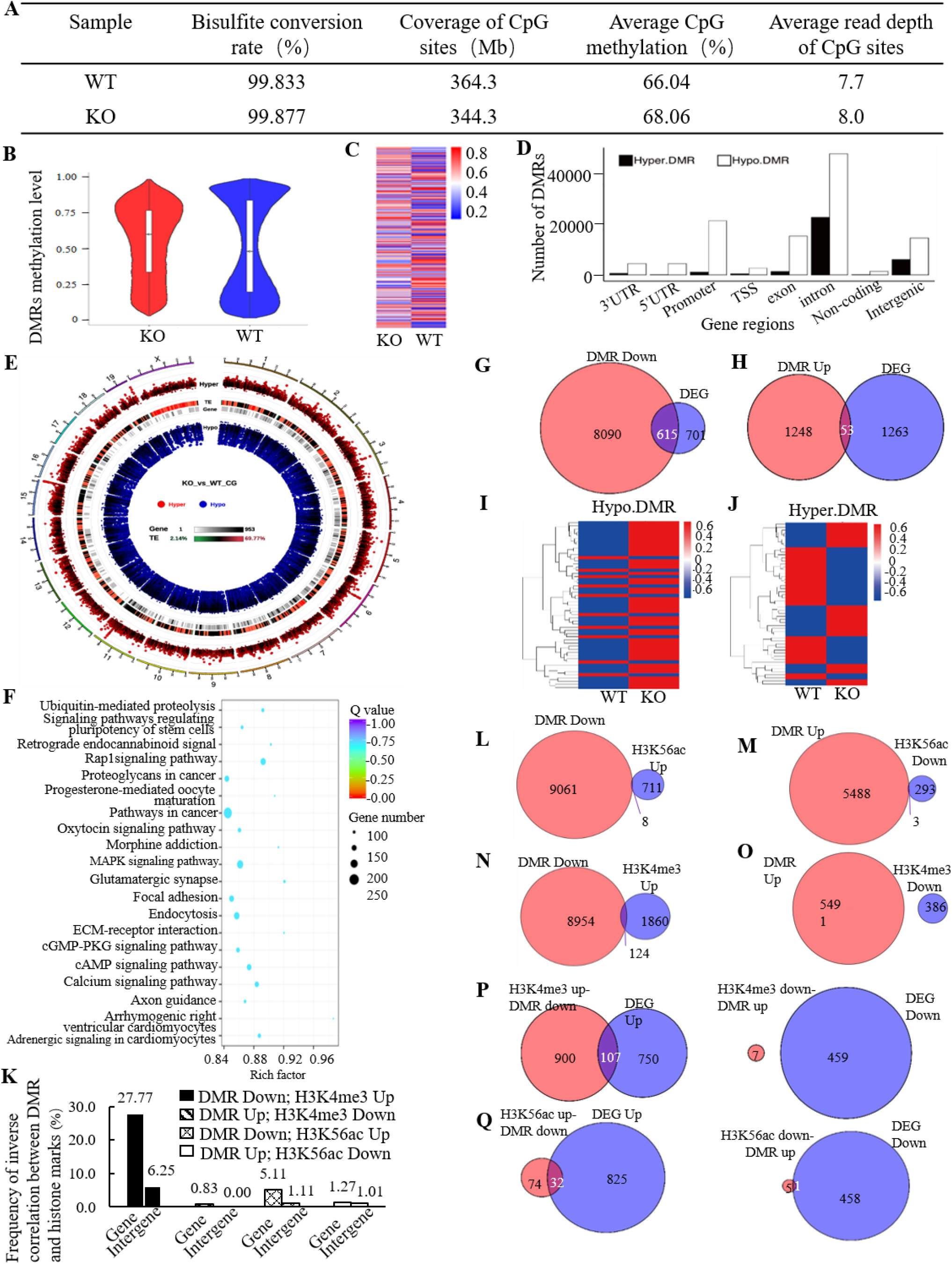
Differential methylome analysis in the wild-type and PA200^−/−^ MEF cells. (*A*) Summary of WGBS analysis from the wild-type (WT) and PA200^−/−^ (KO) MEF cells. (*B*) Violin plots showing the distributions of DNA methylation levels in WT and PA200^−/−^ MEFs. (*C*) Heatmap of methylation levels within CG DMRs in WT and PA200^−/−^ MEF cells. (*D*) Distributions of CG DMRs gene regions in the PA200-deficient MEF cells (normalized to the wild-type group). (*E, F*) Venn plots showing gene numbers of hypo-DMRs (*E*) or hyper-DMRs (*F*) and DEGs in PA200-deficient MEF cells. (*G, H*) Hierarchical clustering of the hypo-DMRs-related DEGs (*G*) or hyper-DMRs-related DEGs (H) in the PA200^+/+^ and PA200^−/−^ MEF cells. (*I*) The frequency of inverse correlation between DMR and peaks of histone marks. Frequency= the peak number of inverse correlation / total peak number of histone marks in gene regions or intergene regions. Gene and intergene regions were defined by the HOMER (v4.9.1). The gene region includes TSS, promoter, intron, and exon. (*J*) (*K*) Venn plots showing peak numbers of H3K56ac and DNA methylation that were upregulated (*J*) or downregulated (*K*) in the intergene regions of the PA200-deficient MEF cells. (*L, M*) Venn plots showing peak numbers of H3K4me3 and DNA methylation that were up-regulated (*L*) or down-regulated (*M*) in the intergene regions of the PA200-deficient MEF cells. (*N*) Venn plots showing numbers of the DEGs and genes that corresponds to the “inverse correlation of H3K4me3 and DMRs” regions in the PA200-deficient MEF cells. (*O*) Venn plots showing numbers of the DEGs and genes that corresponds to the “inverse correlation of H3K56ac and DMRs” regions in the PA200-deficient MEF cells. (*P*) Circular representation of the genome-wide distribution of CG DMRs in the PA200-deficient MEF cells (normalized to the wild-type group). This visualization was generated using the Circos software58. Hyper, hyper-methylated DMRs; Hypo, hypo-methylated DMRs. (*Q*) KEGG pathway enrichment of CG DMRs genes in the PA200-deficient MEF cells (normalized to the wild-type group).

**Fig. S8.**
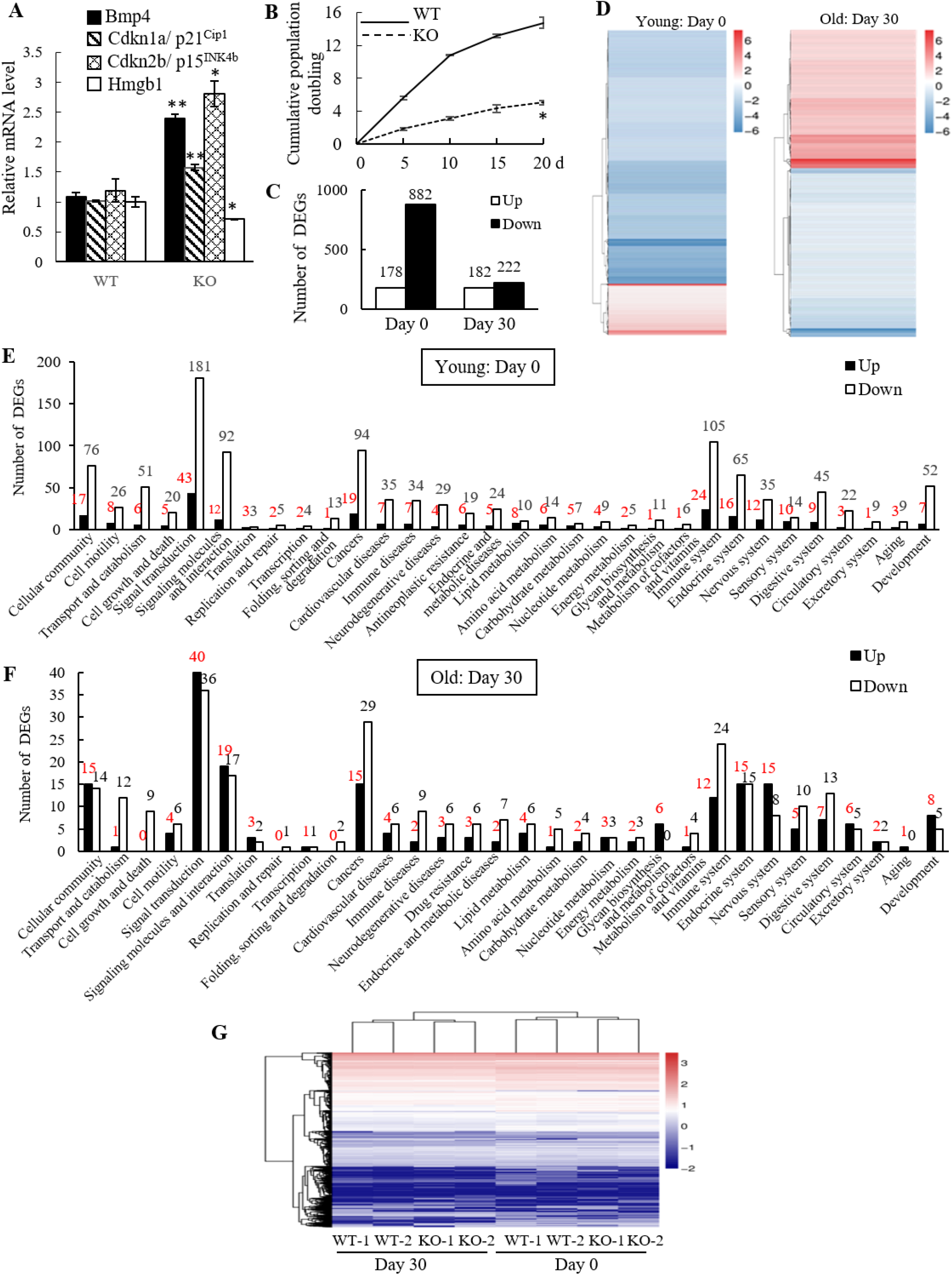
Deletion of PA200 changes gene expression during aging of primary MEF cells. (*A*) Quantitative PCR analysis of 4 selected aging markers in mouse livers. (*B*) Growth curve for the cumulative population doubling of primary wild-type (WT) and PA200-deficient (KO) MEF cells. (*C*) Numbers of the up-regulated and down-regulated genes in the PA200-deficient primary MEF cells (normalized to the wild-type group). (*D*) Hierarchical clustering of intersection DEGs in the PA200-deficient primary MEF cells (normalized to the wild-type group). (*E, F*) KEGG pathway classification of up-regulated and down-regulated DEGs analyzed by RNA-seq in young (*E*) or old (*F)* PA200-deficient primary MEF cells (normalized to the wild-type group). (*G*) Expression heatmap of transcription factor TF genes in the wild-type and PA200-deficient primary MEFs. Coloring indicates the log2 transformed fold change. Data are representative of one experiment with at least three independent biological replicates (mean ± SEM, ** *p*<0.01, * *p*<0.05, two tailed unpaired test).

**Fig. S9.**
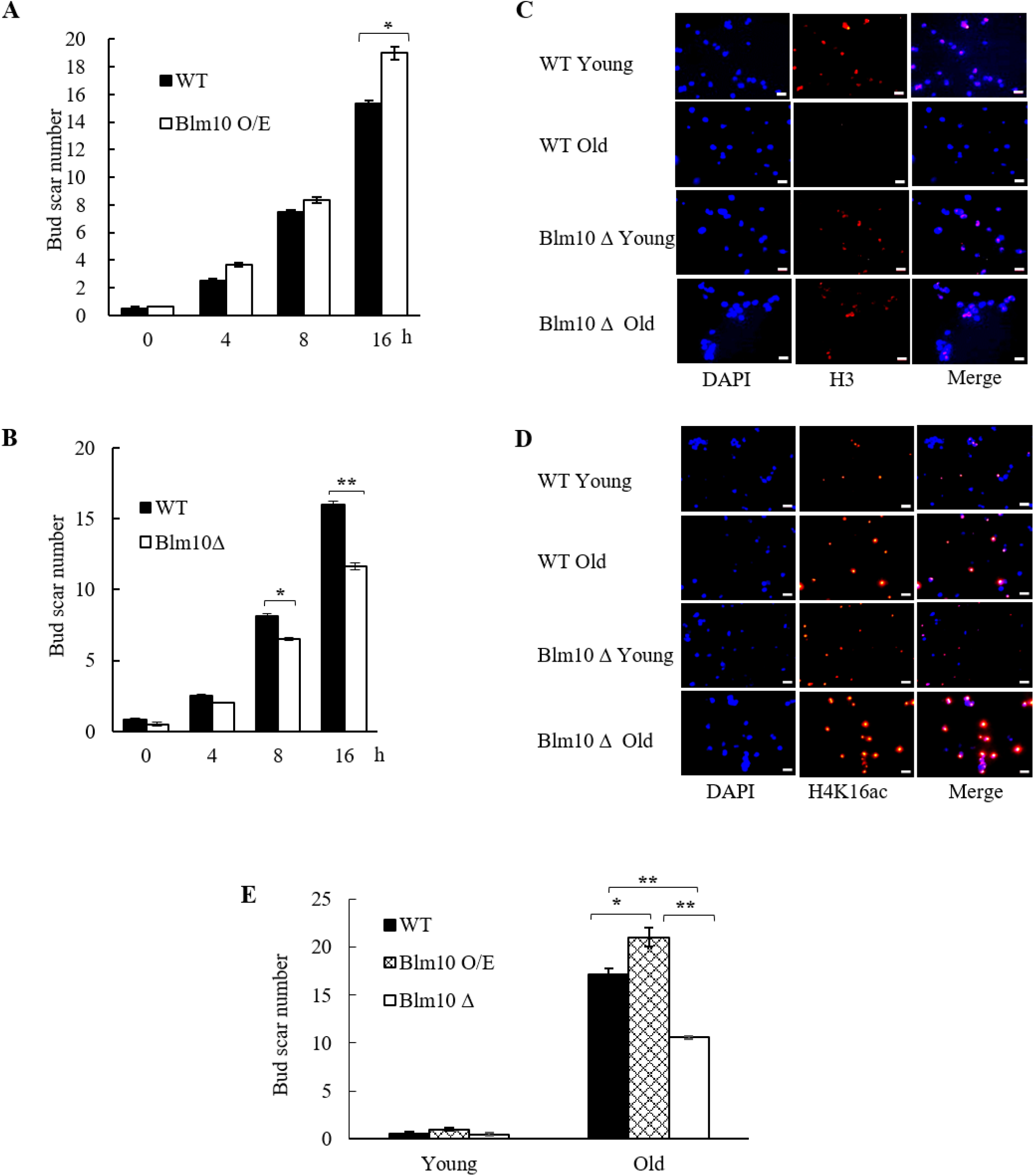
Effects of PA200/Blm10 on the levels of H3, H4 and H4K16ac during cellular aging. (*A, B*) Scar number in the young or old wild-type, Blm10-overexpressing (Fig.5E) and Blm10-deficient (Fig. 5F) yeast. More than 20 cells were measured in each group. (*C, D*) Immunofluorescent images of histone H3 (*C*) and H4K16Ac (*D*) in the young or old wild-type and Blm10-deficient yeast. More than 20 cells were measured in each group. Scale bar, 5 μm. (*E*) Bud scar numbers in yeast as in Fig. 5D. Young: 0 hour; Old: 80 hours. Data are representative of one experiment with two independent biological replicates (mean ± SEM, ** *p* < 0.01, * *p* < 0.05, two tailed unpaired test).

**Table S1.**
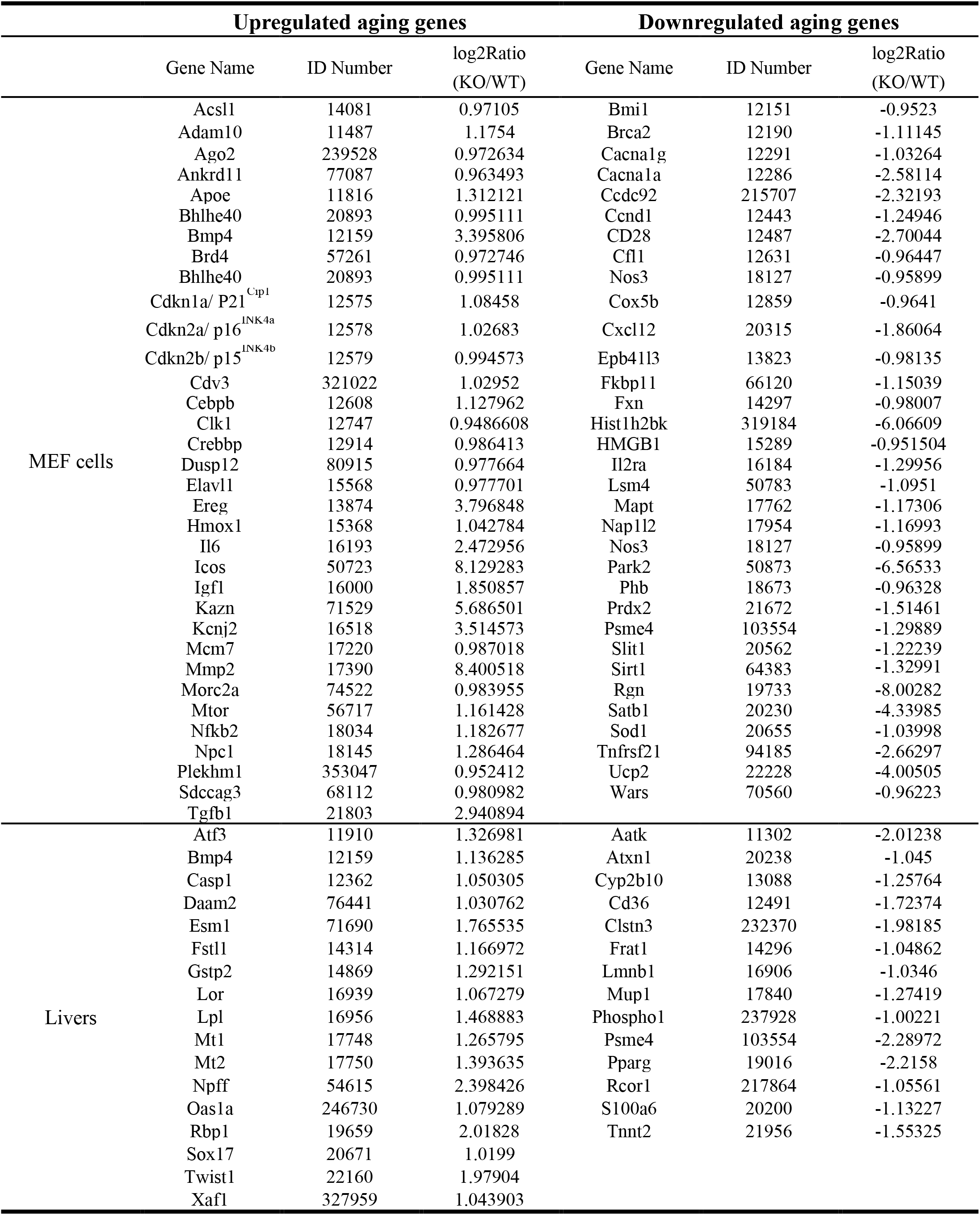
Aging-related DEGs in PA200^−/−^ (KO) and WT MEF cells / Livers.

### Supplementary Methods (SI Methods)

#### Mice

The PA28γ-deficient mice (C57BL/6N) were kindly provided by Drs. Lance Barton and Xiaotao Li. The PA200-deficient mice (C57BL/6N) were generated with the CRISPR/Cas9 technique by Beijing Biocytogen Co., Ltd (Beijing, China). In brief, PA200^−/−^ mice were generated by nonhomology end joining, which was induced by 2 double-strand break repairs after introduction of 2 single-guide RNAs with Cas9. Two single-guide RNAs were designed to target a region upstream of intron 42 and downstream of 3’UTR, respectively. Different concentrations of Cas9 mRNA and single-guide RNAs were mixed and co-injected into the cytoplasm of 1-cell-stage fertilized eggs to generate chimeras. Polymerase chain reaction genotyping and sequencing revealed that some pups carried deletions of about 21-kb spanning 2 single-guide RNA target sites, removing the PA200 amino acid 1606-1843.

For genotyping, DNA was extracted from tip of the tail and analyzed by PCR with the primers: forward primer 5’-TGTTCACCACTGAAGTATAGGAACTCA, reverse primer 5’-GCTGGAGT ATTCTTCCCTTGGGAGT (for PA200^−/−^ mice); forward primer 5’-AGCTGTGAAGATCATG AGCTATAGTAGT, reverse primer 5’-TCTGTGAGTCTGAAGCCTGGTCTAA (for wild-type mice). The animals’ care was in accordance with institutional guidelines.

#### Antibody information

Antibodies against the histone H3 (1:1000, Abcam, #ab1791), H4 (1:3000, EMD Millipore, #05-858), H3.1 (1:1000, Active Motif, #61629), H3.3 (1:1000, Abcam, #ab176840), PA200 (1:500, Abcam, #ab181203), HMGB1 (1:1000, Cell signaling technology, #6893), Lamin B1 (1:5000, Abcam, #ab133741), Rpb1 (1:2000, Abcam,#ab76123), HA (1:2000; Santa Cruze, sc7392), PA28γ (1:3000, Enzo, PW8190), streptavidin-HRP (1:5000, ZSGB-BIO, #ZB-2404), and β-actin (1:5000, Sigma-Aldrich, #A5441) were used as primary antibodies to detect the corresponding proteins. Peroxidase-conjugated anti-mouse IgG (1:5000,ZSGB-BIO,#ZB-5305), anti-rat IgG (1:4000, ZSGB-BIO, #ZB-2307) or anti-rabbit IgG (1:3000, ZSGB-BIO, #ZB-5301) was used as secondary antibody. Antibodies against H3K56ac (Abcam, #ab76307), H3K4me3 (Abcam, #ab8580) and RNA polymerase II (Abcam, #Ab5095) were used for ChIP-seq assay.

#### RNA-seq preparation and data processing

RNA samples were collected from G1-arrested MEF cell lines or mouse livers using TRIZOL reagent, and sequenced on Illumina BGISEQ-500 at Beijing Genomic Institution (BGI, Shenzhen, China; http://www.genomics.org.cn). Clean-tags were aligned to the mm10 reference genome. For gene expression analysis, the matched reads were calculated and then normalized to RPKM using RESM software (1). The significance of the differential expression of genes (DGEs) was defined by the bioinformatics service of BGI according to the combination of the absolute value of log2-Ratio ≥1 and diverge probability≥0.8 (2). Gene Ontology (GO) and pathway annotation and enrichment analyses were based on the Gene Ontology Database (http://www.geneontology.org/) and KEGG pathway database (http://www.genome.jp/kegg/), respectively. The software Cluster and Java Tree view were used for hierarchical cluster analysis of gene expression patterns (3, 4). DEGs were defined according to the combination of the absolute value of log2-Ratio ≥1 and diverge probability≥0.8. Coloring indicates the log2 transformed fold change. We performed hierarchical cluster analysis of DEGs using the heatmap method in R version 3.3.3. We measured the distance between genes using the agglomeration method (Ward.D2).

#### Real-time PCR

The Transcriptor First Strand cDNA synthesis Kit (Roche) was used for cDNA synthesis from total RNA according to manufacturer’s instructions. Quantitative PCR was performed with an ABI 7500 Real-Time PCR System (Applied Biosystems) in technical duplicates from two biological replicates. Gapdh was set as control. Relative expression values were calculated using the ΔCt method. The primer sequences were listed in Table S2.

**Table S2.**
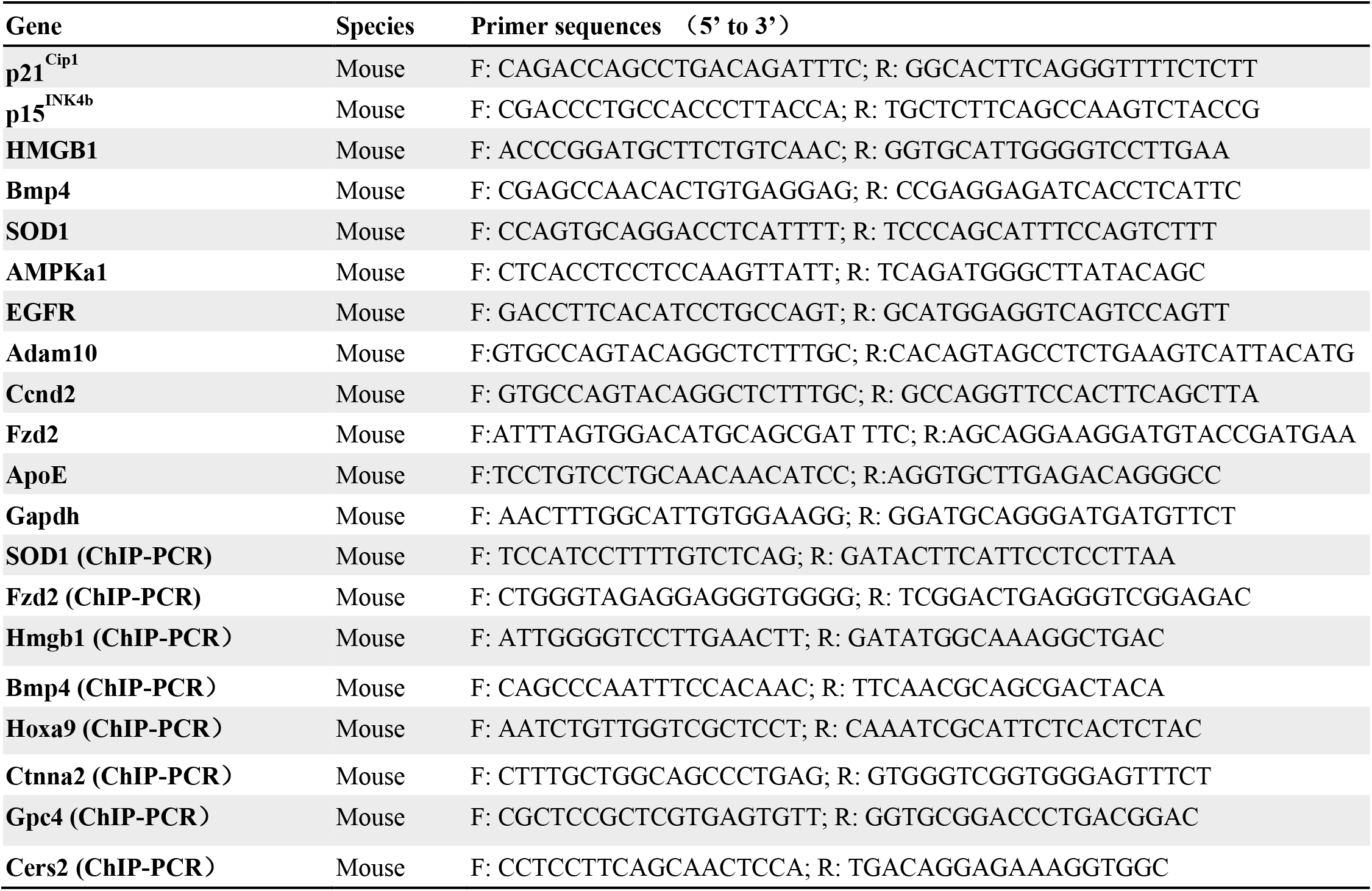
Primers used in this study.

#### ChIP-seq preparation and data processing

About 5 × 10^7^ cells were used for each ChIP-Seq assay. Two micrograms of either histone H3K4me3 antibody or histone H3K56ac antibody were used for each immunoprecipitation reaction. The sequence libraries were generated using the purification kit (CST, #14209) for in-depth whole-genome DNA sequencing by the Illumina HiSeq 2500 platform (BGI, Shenzhen, China; http://www.genomics.org.cn,). ChIP–seq reads were aligned to the mouse reference genome (mm10) using HISAT2 (version 2.0.3). All unmapped reads, non-uniquely mapped reads and PCR duplicates were removed by SAM tools (version 1.3.1). To call peaks, we used MACS2 with the parameters -bdg -broad -nomodel and -SPMR to normalize each sample by sequencing depth, and used subcommand bdgcmp with parameter -m FE to get noise-subtracted tracks. The ChIP–seq signal tracks were visualized in Integrative Genomics Viewer (IGV). To determine the ChIP-seq average profile in genome and in up- or down-regulated genes, we used ngs.plot to calculate mean coverage of all regions, and then used R to visualize. The analyzed functional elements included TSS (transcription start site) and gene body.

#### Analysis of ChIP-seq enrichment patterns

The ChIP-seq average signals at promoters (±2.5 kb) of DEGs were computed by ngs.plot, and gene expression levels (RPKM) were used in the analysis. The correlation between ChIP-seq enrichment at promoters and gene expression was shown in heat maps.

For different ChIP-Seq data (such as H3K4me3 in PA200^−/−^ MEF cells and PA200^+/+^ MEF cells), we used MAnorm to statistically compare the quantitative binding differences and used M values to classify the upregulated or downregulated regions. For the identified up/down-regulated regions of H3K4me3/H3K56ac, we annotated their related genes as enrichment by Homer. The ChIP-seq (polymerase II or histone marks) enrichment patterns at the corresponding genes were calculated by ngs.plot.

#### Whole-genome bisulfite sequencing (WGBS) for DNA methylation

The genome DNA was isolated from MEF cells using TIANamp Genomic DNA Kit (TIANGEN, #DP304) according to manufacturer’s protocol. The purified DNA was then directly proceeded to quality inspection, library construction and sequencing at the Novogene Bioinformatics Institute (Beijing, China) on an Illumina HiSeq 2000/2500 platform. Briefly, approximately 5.2 μg of genomic DNA spiked with 26 ng of lambda DNA was fragmented by sonication to 200-300 bp with Covaris S220, followed by end repair and adenylation. The sonicated DNA was then ligated with cytosine-methylated barcodes. These DNA fragments were treated with bisulfite using EZ DNA Methylation-GoldTM Kit twice. The resulting single-strand DNA fragments were amplified by PCR using KAPA HiFi HotStart Uracil + ReadyMix (2×). Qubit® 2.0 Fluorometern was selected to quantify the library concentration. The insert size was assessed on an Agilent Bioanalyzer 2100 system. Subsequently, 125 bp single-end reads were generated. Image analysis and base calling were performed with Illumina CASAVA pipeline. Finally, 125 bp paired-end reads were generated. Clean reads were obtained as previously described (5). The reference genome was transformed into the bisulfite-converted version (C-to-T and G-to-A converted) and indexed by Bowtie 2. Clean reads were also transformed into the fully bisulfite-converted version and then aligned to the above converted version of the genome. Differentially methylated regions (DMR) were analyzed by DSS software, which is based on beta-binomial distribution, and the related genes of DMRs were annotated. For DMR, we identified the upregulated/hyper-DMR if the mean methylated value of KO sample is larger than that of WT sample, and *vice versa* for the downregulated/hypo-DMR. For analysis of genome-wide methylation, each chromosome was classified into several bins with equal sizes, and the methylation level of each bin was calculated as the proportion (mC counts / mC counts + umC counts). The high methylation level of genome was calculated as the ratio of the number of high-methylated (>0.5) bins to the number of all bins.

KEGG enrichment analysis of DMR-related genes was implemented by the GO-seq R package (6) in which gene length bias was corrected. GO terms with corrected P-value less than 0.05 were considered significantly enriched by DMR-related genes. We used KOBAS software (7) to test the statistical enrichment of DMR-related genes in KEGG pathways.

#### Growth curve assay

Cell population doubling was determined as described (8). Growth rates of primary MEFs were determined by microscopic measurement (OLYMPUS, CKX31) after staining by trypan blue according to the instructions.

#### Senescence-associated β-galactosidase (SA-β-gal) staining assay

Staining was performed using the Senescence Cells Histochemical Staining Kit (Sigma-Aldrich, #CS0030) according to manufacturer’s instructions. Briefly, the cells were seeded at 3 × 105 cells per well in six-well plates for 24 h. The cells were then washed with 1×PBS and fixed with fixation buffer for 7 min at room temperature. The cells were then incubated overnight at 37 °C with the working solution containing 1 mg/ml X-gal in the kit, and senescence was identified as positive in the dark blue-staining cells observed. At least 100 cells were counted for each group in over three random fields to determine the percentage of SA-β-gal-positive cells.

#### 8-arm radial maze

A group of animals were trained so that they would become habituated to the apparatus (Med Associates, ENV-256I) and food pellets for 3 days before each test. The mice were also treated with food limitation to reduce the body weight to 85%. In each training session, the animal was placed in a circular plastic wall on the platform in the middle of the 8-arm radial maze. Then, after 1 min, the ring was lifted and the animal was allowed to move freely in the maze. The trial continued until the animal had either entered all 8 arms or until 10 min had elapsed. A small piece of popcorn (50 mg) was used as the bait. The performance of a given animal in each trial was assessed using three parameters: the number of correct choices in the initial 8 chosen arms, the number of errors which was defined as choosing arms that had already been visited, and the time elapsed before the animal ate all 8 pellets.

#### Measurement of T cell surface marker expression

Spleens from mice were analyzed for the naive/memory CD4 and CD8 T cell subsets by flow cytometry using the following antibodies: CD4 (GK1.5; eBiosciences), CD44 (IM7; eBiosciences), CD62L (MEL-14; eBiosciences). Cells were acquired with a BD Biosciences FACS Aria IIIu instrument and analyzed using the FlowJo (Tree Star, Ashland, OR, USA) software package.

#### Fiber diameter measurements

Fiber diameter measurements were performed on cross sections of gastrocnemius muscles from the wild-type or PA200 deficient mice female mice at 3-month-old or 12-month-old. Gastrocnemius muscles were collected and processed for histology as described previously (9). A total of 50 fibers were measured per muscle using Image J.

#### Determination of glomerular sclerosis

Formalin-fixed and paraffin-embedded kidney samples were stained using routine hematoxylin and eosin. Forty randomly selected glomeruli were scored for sclerosis. Glomeruli with > 50% sclerosis were determined to be sclerotic.

#### Statistical Analysis

Significance levels for comparisons between two groups were determined by the two-tailed unpaired *t*-test, mean ± SEM (*p<0.05 and **p<0.01), normal distribution. All of the images of immunoblotting were chosen blindingly and randomly and quantitated by image J.

## References

1. E. I. Campos, D. Reinberg, Histones: annotating chromatin. Annu Rev Genet 43, 559–599 (2009).

2. A. J. Ruthenburg, C. D. Allis, J. Wysocka, Methylation of lysine 4 on histone H3: intricacy of writing and reading a single epigenetic mark. Mol Cell 25, 15–30 (2007).

3. S. Stejskal et al., Cell cycle-dependent changes in H3K56ac in human cells. Cell Cycle 14, 3851–3863 (2015).

4. B. Zhang et al., Allelic reprogramming of the histone modification H3K4me3 in early mammalian development. Nature 537, 553–557 (2016).

5. J. A. Lessard, G. R. Crabtree, Chromatin regulatory mechanisms in pluripotency. Annu Rev Cell Dev Biol 26, 503–532 (2010).

6. A. Portela, M. Esteller, Epigenetic modifications and human disease. Nat Biotechnol 28, 1057–1068 (2010).

7. S. B. Hake, C. D. Allis, Histone H3 variants and their potential role in indexing mammalian genomes: the “H3 barcode hypothesis”. Proc Natl Acad Sci U S A 103, 6428–6435 (2006).

8. C. Huang, B. Zhu, H3.3 turnover: a mechanism to poise chromatin for transcription, or a response to open chromatin? Bioessays 36, 579–584 (2014).

9. I. Maze et al., Critical Role of Histone Turnover in Neuronal Transcription and Plasticity. Neuron 87, 77–94 (2015).

10. G. A. Collins, A. L. Goldberg, The Logic of the 26S Proteasome. Cell 169, 792–806 (2017).

11. T. X. Jiang, M. Zhao, X. B. Qiu, Substrate receptors of proteasomes. Biol Rev Camb Philos Soc 93, 1765–1777 (2018).

12. A. Navon, A. Ciechanover, The 26 S proteasome: from basic mechanisms to drug targeting. J Biol Chem 284, 33713–33718 (2009).

13. M. Schmidt et al., The HEAT repeat protein Blm10 regulates the yeast proteasome by capping the core particle. Nat Struct Mol Biol 12, 294–303 (2005).

14. V. Ustrell, G. Pratt, C. Gorbea, M. Rechsteiner, Purification and assay of proteasome activator PA200. Methods Enzymol 398, 321–329 (2005).

15. J. Blickwedehl et al., Role for proteasome activator PA200 and postglutamyl proteasome activity in genomic stability. Proc Natl Acad Sci U S A 105, 16165–16170 (2008).

16. M. X. Qian et al., Acetylation-mediated proteasomal degradation of core histones during DNA repair and spermatogenesis. Cell 153, 1012–1024 (2013).

17. I. K. Mandemaker et al., DNA damage-induced replication stress results in PA200-proteasome-mediated degradation of acetylated histones. EMBO Rep 19(2018).

18. M. H. Hauer et al., Histone degradation in response to DNA damage enhances chromatin dynamics and recombination rates. Nat Struct Mol Biol 24, 99–107 (2017).

19. W. Dang et al., Histone H4 lysine 16 acetylation regulates cellular lifespan. Nature 459, 802–807 (2009).

20. J. Feser et al., Elevated histone expression promotes life span extension. Mol Cell 39, 724–735 (2010).

21. V. T. Nguyen et al., In vivo degradation of RNA polymerase II largest subunit triggered by alpha-amanitin. Nucleic Acids Res 24, 2924–2929 (1996).

22. H. Tagami, D. Ray-Gallet, G. Almouzni, Y. Nakatani, Histone H3.1 and H3.3 complexes mediate nucleosome assembly pathways dependent or independent of DNA synthesis. Cell 116, 51–61 (2004).

23. Z. Dou et al., Autophagy mediates degradation of nuclear lamina. Nature 527, 105–109 (2015).

24. A. Kuma et al., The role of autophagy during the early neonatal starvation period. Nature 432, 1032–1036 (2004).

25. D. Shechter, H. L. Dormann, C. D. Allis, S. B. Hake, Extraction, purification and analysis of histones. Nat Protoc 2, 1445–1457 (2007).

26. M. F. Dion et al., Dynamics of replication-independent histone turnover in budding yeast. Science 315, 1405–1408 (2007).

27. C. L. Liu et al., Single-nucleosome mapping of histone modifications in S. cerevisiae. PLoS Biol 3, e328 (2005).

28. E. A. Meyers et al., Increased bone morphogenetic protein signaling contributes to age-related declines in neurogenesis and cognition. Neurobiol Aging 38, 164–175 (2016).

29. J. P. Brown, W. Wei, J. M. Sedivy, Bypass of senescence after disruption of p21CIP1/WAF1 gene in normal diploid human fibroblasts. Science 277, 831–834 (1997).

30. S. Agnihotri, A. Wolf, D. Picard, C. Hawkins, A. Guha, GATA4 is a regulator of astrocyte cell proliferation and apoptosis in the human and murine central nervous system. Oncogene 28, 3033–3046 (2009).

31. M. Collado et al., Inhibition of the phosphoinositide 3-kinase pathway induces a senescence-like arrest mediated by p27Kip1. J Biol Chem 275, 21960–21968 (2000).

32. V. I. Perez et al., Is the oxidative stress theory of aging dead? Biochim Biophys Acta 1790, 1005–1014 (2009).

33. J. Krishnamurthy et al., Ink4a/Arf expression is a biomarker of aging. J Clin Invest 114, 1299–1307 (2004).

34. J. G. Wood et al., Sirtuin activators mimic caloric restriction and delay ageing in metazoans. Nature 430, 686–689 (2004).

35. B. Li, C. N. Dewey, RSEM: accurate transcript quantification from RNA-Seq data with or without a reference genome. BMC Bioinformatics 12, 323 (2011).

36. R. A. Miller, Biomarkers of aging: prediction of longevity by using age-sensitive T-cell subset determinations in a middle-aged, genetically heterogeneous mouse population. J Gerontol A Biol Sci Med Sci 56, B180–186 (2001).

37. R. Schmitt, A. Melk, Molecular mechanisms of renal aging. Kidney Int 92, 569–579 (2017).

38. J. Lexell, K. Henriksson-Larsen, B. Winblad, M. Sjostrom, Distribution of different fiber types in human skeletal muscles: effects of aging studied in whole muscle cross sections. Muscle Nerve 6, 588–595 (1983).

39. B. D. Strahl, C. D. Allis, The language of covalent histone modifications. Nature 403, 41–45 (2000).

40. D. Moazed, Mechanisms for the inheritance of chromatin states. Cell 146, 510–518 (2011).

41. A. Ferdous, T. Kodadek, S. A. Johnston, A nonproteolytic function of the 19S regulatory subunit of the 26S proteasome is required for efficient activated transcription by human RNA polymerase II. Biochemistry 41, 12798–12805 (2002).

42. F. Geng, W. P. Tansey, Similar temporal and spatial recruitment of native 19S and 20S proteasome subunits to transcriptionally active chromatin. Proc Natl Acad Sci U S A 109, 6060–6065 (2012).

43. B. Khor et al., Proteasome activator PA200 is required for normal spermatogenesis. Mol Cell Biol 26, 2999–3007 (2006).

44. J. Borgel et al., Targets and dynamics of promoter DNA methylation during early mouse development. Nat Genet 42, 1093–1100 (2010).

45. C. Lopez-Otin, M. A. Blasco, L. Partridge, M. Serrano, G. Kroemer, The hallmarks of aging. Cell 153, 1194–1217 (2013).

46. L. W. Harries et al., Human aging is characterized by focused changes in gene expression and deregulation of alternative splicing. Aging Cell 10, 868–878 (2011).

47. A. Nicholas et al., Age-related gene-specific changes of A-to-I mRNA editing in the human brain. Mech Ageing Dev 131, 445–447 (2010).

48. J. Campisi, Aging, cellular senescence, and cancer. Annu Rev Physiol 75, 685–705 (2013).

49. N. Loaiza, M. Demaria, Cellular senescence and tumor promotion: Is aging the key? Biochim Biophys Acta 1865, 155–167 (2016).

50. A. V. Lichtenstein, N. P. Kisseljova, DNA methylation and carcinogenesis. Biochemistry (Mosc) 66, 235–255 (2001).

51. D. C. Kraushaar et al., Genome-wide incorporation dynamics reveal distinct categories of turnover for the histone variant H3.3. Genome Biol 14, R121 (2013).

52. R. B. Deal, J. G. Henikoff, S. Henikoff, Genome-wide kinetics of nucleosome turnover determined by metabolic labeling of histones. Science 328, 1161–1164 (2010).

53. T. Smeal, J. Claus, B. Kennedy, F. Cole, L. Guarente, Loss of transcriptional silencing causes sterility in old mother cells of S. cerevisiae. Cell 84, 633–642 (1996).

## Supplementary References

1. B. Li, C. N. Dewey, RSEM: accurate transcript quantification from RNA-Seq data with or without a reference genome. BMC Bioinformatics 12, 323 (2011).

2. S. Tarazona et al., Data quality aware analysis of differential expression in RNA-seq with NOISeq R/Bioc package. Nucleic Acids Res 43, e140 (2015).

3. M. J. de Hoon, S. Imoto, J. Nolan, S. Miyano, Open source clustering software. Bioinformatics 20, 1453–1454 (2004).

4. A. J. Saldanha, Java Treeview--extensible visualization of microarray data. Bioinformatics 20, 3246–3248 (2004).

5. S. Zhang et al., Genome-wide analysis of DNA methylation profiles in a senescence-accelerated mouse prone 8 brain using whole-genome bisulfite sequencing. Bioinformatics 33, 1591–1595 (2017).

6. M. D. Young, M. J. Wakefield, G. K. Smyth, A. Oshlack, Gene ontology analysis for RNA-seq: accounting for selection bias. Genome Biol 11, R14 (2010).

7. X. Mao, T. Cai, J. G. Olyarchuk, L. Wei, Automated genome annotation and pathway identification using the KEGG Orthology (KO) as a controlled vocabulary. Bioinformatics 21, 3787–3793 (2005).

8. G. H. Liu et al., Progressive degeneration of human neural stem cells caused by pathogenic LRRK2. Nature 491, 603–607 (2012).

9. D. J. Baker et al., Opposing roles for p16Ink4a and p19Arf in senescence and ageing caused by BubR1 insufficiency. Nat Cell Biol 10, 825–836 (2008).

